# MoE-Bind: Guiding De Novo Protein Binder Generation with Sparse Experts

**DOI:** 10.64898/2026.06.13.732043

**Authors:** Dipayan Sarkar, Chiranjib Sarkar

## Abstract

De novo protein binder design has been dominated by structure-based pipelines that require known three-dimensional target conformations and consume substantial compute and generation time per design, limiting their throughput and accessibility for routine large-scale binder exploration. Sequence-only generative models promise a faster and lighter alternative, yet existing systems remain uniformly dense and frequently reintroduce structural computation at inference, undermining the core advantages they were intended to deliver. Across the broader language modelling community, transformers have meanwhile transitioned from fully dense designs to sparse Mixture-of-Experts architectures that decouple capacity from per-token compute, a shift that has yet to reach sequence-only protein binder generation. We present MoE-Bind, an autoregressive protein binder generator that, for the first time in this domain, combines Multi-head Latent Attention with a sparse Mixture-of-Experts feed-forward network and is evaluated under two independent structure predictors, Boltz-2 and AlphaFold2-Multimer. Despite activating less than half the per-token parameters of compute-matched dense baselines, MoE-Bind matches or exceeds them on full-length receptor-conditioned binder generation on a leakage-free Docking Benchmark 5.0 evaluation, transfers without peptide-specific training to short-peptide design, and reduces training and inference compute by a large margin. Routing analysis on generated binders reveals interpretable expert specialization at both the individual amino acid and biochemical group level, a structured expert–token alignment not previously reported for natural-language MoE models. These results show that sparse architectural design, rather than scale, can deliver fast, structure-free, and interpretable protein binder generation.

## 1 Introduction

Designing a protein that selectively binds a chosen target sequence is one of the central open problems of computational biology, sitting at the heart of nearly every therapeutic and biotechnological pipeline that aims to decipher protein–protein interactions (PPIs), from monoclonal antibodies and vaccine immunogens to enzyme inhibitors and modulators of disease-associated interactomes [1–4]. The space of possible protein sequences is astronomically large, and only a tiny fraction of it yields a sequence that folds, stays stable, and binds a chosen target well. To search this space, the field has long relied on experimental methods such as phage display, directed evolution, and high-throughput screening. These methods remain slow, resource-intensive, and largely restricted to targets that are stable and structurally well characterized [5,6], leaving most of the proteome, including intrinsically disordered, conformationally flexible, and poorly resolved targets, effectively out of reach. The result is a sustained demand for computational frameworks that can propose candidate binders de novo from minimal target information, at a scale and turnaround time compatible with modern therapeutic discovery pipelines.

A dominant line of computational binder design has framed the problem as fundamentally structure-centric, in which a high-resolution model of the target together with explicit specification of the binding interface drives a combinatorial search over backbones and side chains compatible with the desired interaction [1, 6]. The infusion of deep learning into this pipeline, through accurate complex prediction [7, 8], structure-aware inverse sequence design [9], and more recently generative diffusion-based models [10–15], has delivered binders with experimentally validated picomolar affinities and now sets the state of the art for de novo binder design [16]. These successes come with substantial operational costs. Each design typically requires repeated structure prediction and inverse-folding passes, so generating and filtering the thousands of candidates needed to identify a small number of viable binders demands tens of GPU-hours per target on high-end accelerators, with the cost growing sharply for longer binders and larger receptors that exceed the memory of commodity hardware. The practical consequence is that high-throughput structure-based binder design at full scale remains accessible mainly to well-resourced industrial and academic groups, and is difficult to deploy in settings where many targets must be explored quickly or where local compute is limited. A lighter, faster generation step that operates directly on protein sequences, without invoking any explicit structural representation, would lower this barrier considerably, and could either stand alone as a binder generator or feed a small, high-quality candidate pool into subsequent structure-based refinement.

The foundation for such a sequence-only approach has already been laid by the protein language modelling community, which has established that self-supervised training on unaligned amino acid sequences alone is sufficient to recover deep biological signal, including secondary structure, residue–residue contacts, function, and, at sufficient scale, atomic-level three-dimensional geometry [17–22]. Building on this foundation, binder generation is increasingly being framed as a conditional sequence-modelling task, in which an autoregressive decoder produces a candidate binder one residue at a time given the target sequence as a prompt, without ever invoking an explicit structural representation of either partner. This formulation addresses the operational bottlenecks of structure-based design directly. It generalises to any target with a known primary sequence, sidesteps the combinatorial expense of structural sampling, and, by avoiding repeated full-atom structure prediction in the generation loop, can produce hundreds of candidate binders per target within seconds rather than the hours required by structure-dependent pipelines.

Despite this promise, existing sequence-only protein binder generators are uniformly dense transformers, even as the broader large language model landscape has shifted decisively toward sparse Mixture-of-Experts (MoE) architectures [23–25]. For protein sequence modelling this gap is particularly consequential, because the motivation for Mixture of Expert (MoE) is structural rather than incidental. The feed-forward sub-layer accounts for roughly two-thirds of the parameters of a standard transformer block [26], a share that rises further when modern compressed-attention schemes such as MLA are used in place of standard multi-head attention [24], so the feed-forward stack is where almost all of a protein language model’s representational capacity for learning amino acid statistics, residue co-occurrence patterns, and target–binder interaction signals actually resides. Sparsifying this stack by replacing it with a routed bank of experts therefore yields the largest possible decoupling of total parameter count from per-token compute, while leaving the attention pathway, which already handles long-range receptor context efficiently in modern protein decoders, untouched. In a MoE layer, each input residue token is routed through a learned gating function to a small subset of specialised expert feed-forward sub-networks rather than through a single shared stack, so additional capacity can be added by introducing more experts without proportionally increasing the active parameters or floating-point operations consumed per generated residue, and explicit room is created for different experts to specialise in different types of biological input [25]. Because the feed-forward layer is the site at which most of the model’s biological knowledge is encoded, making it the locus of specialisation is precisely what unlocks the scaling and efficiency advantages of sparse models for protein binder generation, where every receptor query benefits from the targeted activation of only the experts whose learned residue-level patterns are relevant to that specific target context.

In natural language, this promise of expert specialisation is realised only diffusely. The token vocabulary spans tens of thousands of sub-word units, and expert specialisation, while measurable, tends to be distributed and difficult to interpret in terms of any single linguistic category [23]. Proteins occupy the opposite end of the spectrum, with an effective vocabulary of only twenty canonical amino acids that fall into a small number of well-characterised biochemical groups such as hydrophobic, polar, charged, and aromatic residues. This tiny, biologically structured token alphabet creates a uniquely favourable regime for MoE specialisation, in which individual experts could plausibly align with individual residues or with entire biochemical classes, and in which routing decisions can be inspected at residue resolution rather than being buried inside a large sub-word vocabulary. Whether such residue-level or group-level expert specialisation actually emerges in a generative protein model has, to the best of our knowledge, never been examined. Prior MoE protein systems have either focused on representation learning rather than target-conditioned generation [27], or restricted their analysis of routing to antibody-specific encoders without inspecting per-residue or per-biochemical-group expert allocation [28], leaving open the more fundamental question of how a sparse generative protein decoder organises its twenty-amino-acid alphabet across specialised experts. Equally unaddressed is whether any such specialisation, if it emerges, translates into measurable gains for target-conditioned binder design, where the routing pattern of every generated residue can in principle be tied back to the biochemical identity of that residue and to the receptor context that produced it. The present work is, to our knowledge, the first to directly probe residue-resolved expert behaviour in a sparse generative protein binder model and to connect it to downstream binder quality.

Sequence-only binder generation also benefits from advances in attention design that target the memory and latency bottlenecks of decoder-only generation, both of which are sharpened by the unusual length distribution of protein sequences. Receptors are often several hundred residues long and must be cached in full while the binder is generated residue by residue, so attention memory grows quickly with target length. Multi-head Latent Attention (MLA) [24] addresses this by compressing the representation that is cached for each attention head, projecting keys and values into a low-rank latent space and thereby decoupling the cached state from the number of attention heads. The result is a dramatic reduction in key-value cache memory at inference, which directly matters for protein binder generation, where hundreds of candidate sequences must typically be sampled for every receptor and each sample carries the full receptor context in cache. Unlike multi-query and grouped-query attention schemes [29], which save memory by tying or sharing heads and therefore degrade the per-residue attention signal that protein decoders depend on to align binders to specific receptor positions, MLA preserves full per-head expressivity while still compressing the cache. MLA and MoE therefore tackle the two complementary bottlenecks of sequence-only binder generation, active compute in the feed-forward stack and cached receptor context in the attention stack, together enabling fast, lightweight, target-conditioned generation even for long therapeutic receptors.

Motivated by these gaps, we present MoE-Bind, a sequence-only generative model for de novo protein binder design built around an MLA-MoE autoregressive decoder-only transformer. MoE-Bind is pre-trained on UniRef50 [30] to internalize general protein sequence statistics, then instruction fine-tuned on a redundancy-filtered subset of high-confidence physical interactions from STRING v12 [31, 32], and evaluated on held-out complexes drawn from the Protein-Protein Docking Benchmark 5.0 [33] and the peptide-binder test set [34, 35]. The design loop itself remains fully structure-free, with structure used only as an external sanity check on generated binders, co-folded with Boltz-2 [36,37] and independently with AlphaFold2-Multimer via ColabFold [7, 8] and scored by interface predicted TM-score, so that reported gains reflect genuine binder quality rather than artifacts of any single predictor. At matched compute, the 100M-parameter MoE-Bind model meets or exceeds the structural hit rate of compute-matched dense MHA and GQA baselines of the same nominal scale while activating less than half of their per-token parameters, demonstrating that the sparse MLA-MoE design delivers a clean compute-quality advantage at small scale rather than only at the multi-billion parameter regime characteristic of natural-language MoE systems. Detailed routing analysis on generated binders further reveals that MoE experts specialize both at the level of individual amino acids and at the level of biochemical groups, a degree of interpretable specialization that has not, to the best of our knowledge, been reported for natural-language MoE models, and that we attribute to the unusually small and biochemically structured protein token alphabet. This residue-resolved specialization opens a direction for future work in which experts can be inspected, pruned, or specialised by biochemical class to inject prior biological knowledge into sequence-only binder generators. Beyond the architectural and interpretability contributions, the combination of MLA-driven compression and MoE-driven sparse activation enables hundreds of full-length binder candidates to be generated per target in seconds on a single accelerator, restoring the speed and accessibility advantages that motivate structure-free generation in the first place.

## 2 Related Work

Computational protein binder design has been dominated by structure-dependent methods such as RFDiffusion [11] and AlphaProteo [12], which require high-resolution target structures and explicit binding-site specifications. Sequence-only approaches have emerged as structure-free alternatives. PPI-LLaMA2 [38] fine-tunes a 342M-parameter LLaMA2 decoder [39] directly on PPI pairs from IntACT and BioGRID, autoregressively decoding binder sequences conditioned on a receptor prompt via a special ”separate” delimiter token, and demonstrates competitive AlphaFold2-Multimer ipTM scores against RFDiffusion on 15 DB5 targets, without any structural input. Prot42 [40] scales this paradigm to 500M to 1.1B parameters by first pre-training on 57.1M UniRef50 sequences and then instruction fine-tuning on 74,000 STRING PPI pairs, extending the context window to 8,192 tokens to capture long-range dependencies; it achieves strong in silico binding affinities on therapeutic targets including VEGF-A and SARS-CoV-2 RBD. Both methods, however, carry limitations that motivate the present work.PPI-LLaMA2 employs a BPE tokenizer rather than the character-level amino acid tokenization standard in protein language modelling [17, 18], which may obscure per-residue signals at binding interfaces; its 342M-parameter dense model also trains without protein-sequence pre-training, potentially limiting what the model can learn from a small interaction-curated corpus, and a small evaluation process leaves generalisation across protein families largely uncharacterised. Prot42, while benefiting from UniRef50 pre-training, fine-tunes on sequences capped at ≤250 amino acids per chain with a combined target-binder context of only 512 tokens, which is well below the 8,192-token pre-training window its architecture supports, suggesting that the model’s generative capability may not transfer to the longer, multi-domain receptors that characterise most relevant targets. At inference, Prot42 generates 500 candidate sequences per target and then runs Boltz-1 structure prediction on each to retain only those within 6–8Å of pre-specified binding hotspot residues; this pipeline not only requires prior knowledge of interface residues but also incurs the cost of 500 structure predictions per target, likely impacting negatively the speed and accessibility advantages that sequence-only binder generation is intended to provide over structure-dependent methods. On the peptide design front, PepMLM [34] fine-tunes ESM-2 [18] via a masking strategy that reconstructs short peptides (≤50 residues) appended to the target C-terminus, achieving a 38% ipTM hit rate with experimental validation on NCAM1 and disease-relevant targets; however, this encoder-only formulation strictly specific for short peptides of length ≤50 residues. In the MoE domain, AIDO-Protein [27] introduces the first sparse MoE protein language model of 16B total parameters, top-2 routing over 8 experts, pre-trained on 1.2T amino acids, achieving state-of-the-art results across the xTrimoPGLM benchmark and demonstrating that sparse architectures can outperform dense counterparts at equivalent active compute; it is a transformer encoder-only model trained with a masked language modelling objective and, while adapted for structure-conditioned sequence generation (inverse folding), it has no target-sequence-conditioned generative capability for binder design. Similarly, BALM-MoE [28] applies sparse Top-2 MoE routing to antibody language models (200M active / 710M total), showing that token-choice routing yields expert specialisation in diverse CDRH3 regions , but remains an antibody-specific encoder focused on representation rather than generation. On the accessibility front, none of the models provides an official web server for binder generation, both are distributed as GitHub code repositories, with moPPIt restricted to 3 an academic-only non-commercial licence and accessible only via Colab notebooks. Beyond accessibility, both tools are limited to short linear peptides (≤50 residues) and rely on AlphaFold2-Multimer or ESMFold to evaluate candidates, reintroducing structural computation into an ostensibly sequence-only pipeline.

Collectively, these works expose three gaps in the current landscape of computational protein binder design. State-of-the-art structure-based methods such as RFdiffusion and AlphaProteo deliver high success rates but require known three-dimensional target conformations and consume substantial compute per design, restricting their accessibility to well-resourced laboratories. Sequence-based generative models for protein binder design have, despite the decisive shift of large language models toward sparse Mixture-of-Experts architectures [24,25] for their well-documented advantages over dense transformers [29, 41], remained uniformly dense, leaving the combination of Multi-head Latent Attention with sparse Mixture-of-Experts routing unexplored in the binder design domain. The most recent sequence-only pipelines, in turn, reintroduce structural dependence at inference time, with Prot42 filtering candidates via Boltz-1 structure prediction and PepMLM ranking outputs using ESMFold, undermining the speed and accessibility advantages that motivate structure-free generation in the first place. MOE-Bind directly answers these gaps by design rather than by scale. The dependence on three-dimensional target conformations is removed by conditioning generation on the receptor amino acid sequence alone. The architectural homogeneity of prior sequence-only binder generators is replaced by an MLA-MoE backbone, the first time, to the best of our knowledge, that latent-attention compression and sparse mixture-of-experts routing are combined in a target-conditioned protein binder generator. The inference-time structural dependence of Prot42 and PepMLM is avoided by ranking generated binders with sequence-intrinsic measures, with full-atom structure prediction reserved exclusively for external evaluation under Boltz-2 and AlphaFold2-Multimer rather than for generated binder selection.

## 3 Materials and Methods

### 3.1 Problem Formulation

We treat the protein binder generation as a conditional sequence generation problem. The model receives the target protein sequence as input and generates the binder sequence autoregressively, where each token is an amino acid residue. The model is trained to predict each binder residue given all previous target sequence tokens and the previously generated binder context. During training, the loss is computed over both the target and the binder tokens, so the model learns the sequence architecture of the target and binder together to update its parameters. This approach allows the model to learn the joint distribution of target and binder sequences, potentially improving its ability to generate binders that fundamentally more aligned to bind with the target. Given a target protein sequence *X* = (*x*_1_*, x*_2_*, . . . , x_m_*) of length *m*, the objective is to generate a binder sequence *Y* = (*y*_1_*, y*_2_*, . . . , y_n_*) of length *n*, where each *x_i_, y_j_* ∈ V and V is the amino acid vocabulary of 20 standard residues plus special tokens.The model operates without any structural information, relying purely on amino acid sequences, making it applicable to any target with a known sequence regardless of whether an experimentally resolved structure exists.

### 3.2 Dataset Construction

Three types of datasets were used in this study, including a large corpus of protein sequences for pre-training, a carefully curated set of target-binder protein sequence pairs for fine-tuning, and a separate benchmarking dataset of target proteins for evaluation. The overall representation of the dataset is available in Table 1

**Table 1:** Datasets used in this study and the amount of data retained after filtering. Pre-training and fine-tuning corpora are reported as sequences/pairs; benchmark sets are reported as complexes (targets).

| Dataset | Source | Filtering | Final size |
| --- | --- | --- | --- |
| *Pre-training* |  |  |  |
| UniRef50 | UniProt | Redundancy and length | $\approx 2\text{B}$ tokens<br>(used 200M and 76M tokens) |
| *Fine-tuning* |  |  |  |
| STRING v12.0 | Physical PPI, conf. $\geq 900$ | Redundancy and length | $\approx 2.1\text{M}$ pairs |
| *Benchmarking* |  |  |  |
| DB5 (primary) | DB5, 230 complexes | Length and strict homology | 22 targets |
| DB5 (larger) | DB5, 230 complexes | Length and Relaxed homology | 78 targets |
| Propedia dataset | 203 receptor-peptide | Overlap removal vs. fine-tuning data | 68 pairs |

**Pre-training dataset.** Pre-training sequences were sourced from UniRef50, a non-redundant clustering of the UniProt data in which all sequences within a cluster share at least 50% pairwise sequence identity and only the cluster representative sequence is retained [30]. UniRef50 was chosen as the pre-training dataset by ESM-2 [18], Prot-GPT2 [17] and ProtTrans [19], because it provides broad taxonomic coverage of known protein sequence space while suppressing redundancy, making it well-suited for learning general protein sequence statistics. Apart from a sequence-length cap of 768 residues, no additional filtering beyond the UniRef50 clustering itself was applied, as redundancy reduction is already built into the UniRef50 construction procedure. The corpus was tokenized into approximately 2 billion tokens (a 98:2 train/validation split). Under the 10% Chinchilla optimal token budget, each model consumes only a fraction of this corpus during training, 200M tokens (100M tier) and 76 M tokens (38M tier).

**Finetuning dataset.** Finetuning data was derived from the STRING database (v12.0), which integrates functional and physical protein-protein interaction (PPI) evidence from multiple organisms and data sources [31]. STRING assigns a combined confidence score (0 to 1000) to each interaction by integrating evidence from co-expression, co-occurrence, text mining, and experimental sources. We used the physical interaction subset with only A-B pairs, retaining only interactions with a combined confidence score of at least 900, the highest confidence tier in STRING, obtained by direct experimental evidence. The raw STRING physical interaction set (confidence *>* 900) contained approximately 33 million pairs, heavily redundant due to homologous proteins across organisms. All unique protein sequences were clustered using MMseqs2 [32] at 40% minimum sequence identity with 80% bidirectional coverage. This reduced the dataset from 33 million to approximately 3.1 million non-redundant pairs which is a 90% reduction. After additionally discarding pairs whose combined length less than 30 or exceeding the model’s 768 token context window 2,099,586 usable pairs remained.

**Benchmarking dataset.** Evaluation of generated binders requires a set of protein-protein complexes with experimentally resolved three-dimensional structures. We used the Protein-Protein Docking Benchmark 5.0 (DB5) [33], a curated collection of 230 non-redundant protein complexes with known PDB structures, widely used as a standard benchmark for protein docking and interface prediction methods. From each DB5 complex, the receptor and ligand chain sequences were extracted. Pairs were filtered by length (minimum 30 residues per chain; maximum combined length of 768 residues) to remain within the model’s context window. To prevent any overlap between the benchmark and the pre-training corpus, each DB5 protein was checked for homology against the UniRef50 pre-training sequences and STRING filtered finetuning dataset using MMseqs2 [32]. Any

DB5 protein with sequence identity ≥ 10% at ≥ 80% coverage to any sequence in the pre-training set or finetuning set was removed from the benchmark dataset. This aggressive homology cutoff ensures that benchmark targets are genuinely unseen during pre-training or finetuning, providing an unbiased evaluation of the model’s generalization to novel PPI complexes. The final filtered DB5 set contains 22 complexes that passed both length and homology criteria and constitutes the held-out benchmark the generated binders. To test whether our findings are robust to benchmark size, we constructed a larger held-out set from the same 230 DB5 complexes under a less aggressive, fine-tuning-only leakage filter, receptor and ligand sequences were screened against the STRING physical-interaction fine-tuning corpus with MMseqs2 [32] at 40% sequence identity and 80% bidirectional coverage. Removing 91 complexes for homology and 18 for exceeding the 768-token limit left 121 leakage-free pairs. For the Boltz-2 structural evaluation this set was further restricted to targets for which the native reference and all five models produced a successful Boltz-2 MSA prediction (82 fully paired targets), and three groups of targets sharing identical receptor and binder sequences under different identifiers were each collapsed to a single entry (using the mean reference ipTM as the hit threshold and each model’s lowest-perplexity binder as its representative), yielding the final 78-target benchmark on which all Boltz-2 results are reported. To situate our models relative to a dedicated peptide-binder baseline, we additionally evaluated against the PepMLM test set [34], a curated collection of 203 receptor–peptide complexes of gold standard Propedia dataset [35, 42] with short binders (4–50 residues) drawn from the PDB and used in the original PepMLM-650M evaluation. Because our fine-tuning corpus is derived from STRING DB, a substantial fraction of these receptors overlaps with sequences seen during finetuning. We therefore screened all 203 PepMLM test receptors against the STRING fine-tuning sequences using MMseqs2 with the same 80% sequence identity threshold used in PepMLM’s own dataset, leaving 68 uncontaminated receptor–peptide pairs that constitute our PepMLM comparison benchmark. This filtering ensures that the PepMLM comparison evaluates genuine out-of-distribution generalization rather than memorized fine-tuning targets.

### 3.3 Model Architectures

For the experiments we compare three attention mechanisms, each instantiated in a representative decoder-only causal language model trained with next-token prediction on protein sequences. All three share the same generation pipeline (Figure 1),the vocabulary consists of 31 tokens (20 standard amino acids, special structural delimiters, and control tokens). All models use a maximum context length of 768 tokens, and parameter budgets are matched across architectures at two scales (100M and 38M total parameters) to enable fair architectural comparison. The complete architectural hyperparameters for all three models are detailed in Table 2.

**Figure 1:**
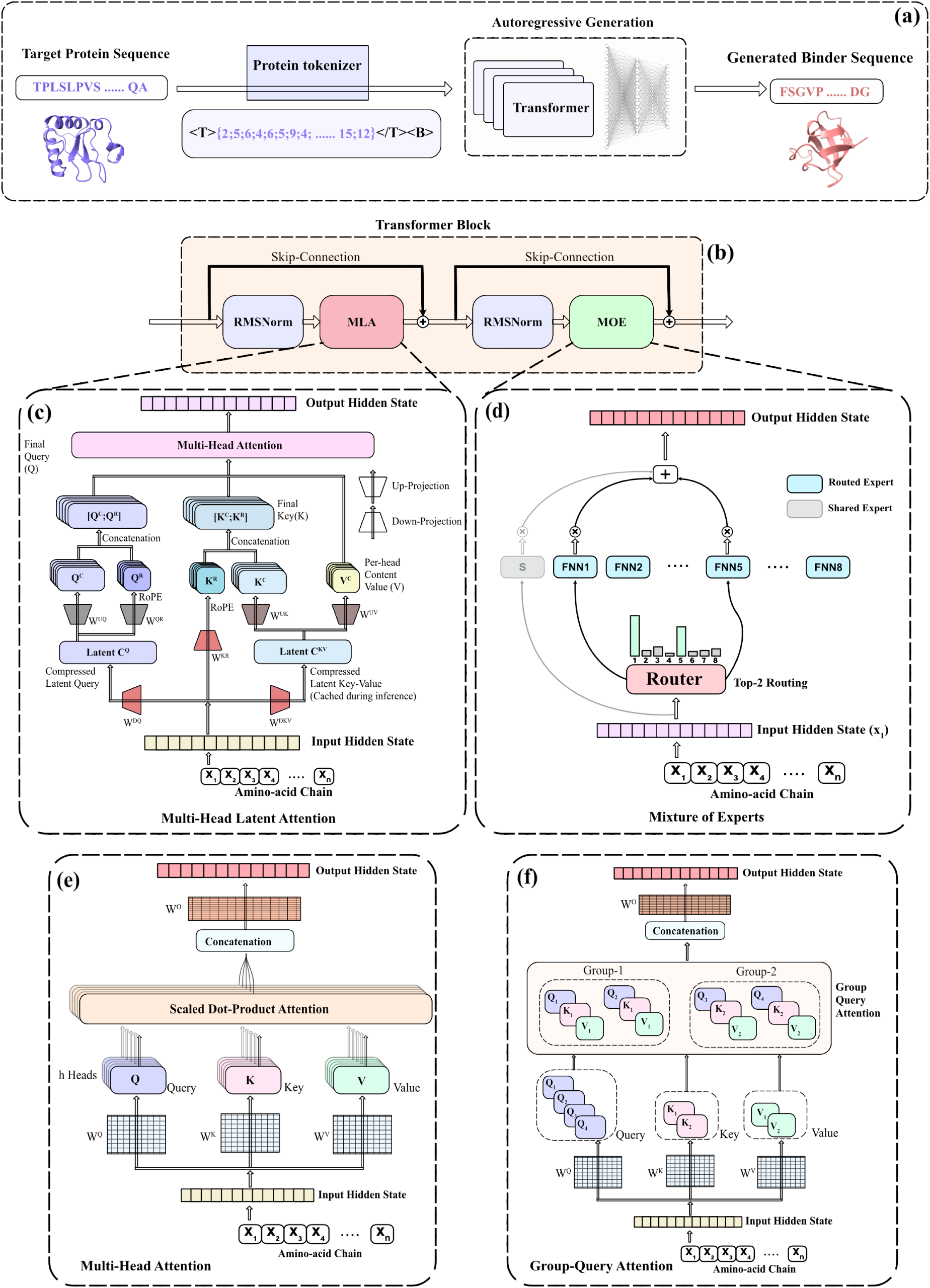
Overview of sequence conditioned the binder generation and model architectures: (a) The sequence conditioned autoregressive binder generation pipeline; (b) the MoE-Bind transformer block (RMSNorm, MLA, MoE with skip connections);(c) Multi-Head Latent Attention with latent compression and decoupled RoPE, (d) Mixture-of-Experts with a top 2 routing and a shared expert, (e) dense Multi-Head Attention, and (f) Grouped-Query Attention.

**Table 2:** Complete architectural hyperparameters for all five models. ^†^ Effective attention dimension per head is 96 for MoE-Bind (*d*_head_ + *r*_rope_ = 64 + 32, per-head dimension *d*_head_ = *d/H*).

|  | MHA-100M | MHA-38M | GQA-100M | GQA-38M | MoE-Bind-100M |
| --- | --- | --- | --- | --- | --- |
| *Architecture* |  |  |  |  |  |
| Attention mechanism | MHA | MHA | GQA | GQA | MLA |
| FFN type | GELU (dense) | GELU (dense) | SwiGLU (dense) | SwiGLU (dense) | SwiGLU (MoE) |
| Normalisation | LayerNorm | LayerNorm | RMSNorm | RMSNorm | RMSNorm |
| Position encoding | Learned abs. | Learned abs. | RoPE | RoPE | RoPE (decoupled) |
| Bias in linear layers | Yes | Yes | No | No | Yes |
| *Model Dimensions* |  |  |  |  |  |
| Embedding dim ($d$) | 768 | 512 | 768 | 512 | 512 |
| Layers ($L$) | 14 | 12 | 16 | 12 | 12 |
| Attention heads ($H$) | 12 | 8 | 12 | 8 | 8 |
| KV heads ($G$) | 12 (MHA) | 8 (MHA) | 4 | 4 | (MLA) |
| Head dim ($d_{\text{head}}$) | 64 | 64 | 64 | 64 | 64<sup>†</sup> |
| KV repetition ($H/G$) | 1× | 1× | 3× | 2× | (MLA) |
| FFN hidden dim ($h$) | 3,072 | 2,048 | 2,048 | 1,536 | 576 (per expert) |
| FFN expansion ($h/d$) | 4.0× | 4.0× | 2.67× | 3.0× | 1.125× |
| *MLA Configuration* |  |  |  |  |  |
| KV LoRA rank ($r_{KV}$) | — | — | — | — | 64 |
| Q LoRA rank ($r_Q$) | — | — | — | — | 64 |
| RoPE dim per head ($r_{\text{rope}}$) | — | — | — | — | 32 |
| *MoE Configuration* |  |  |  |  |  |
| Total routed experts ($N$) | — | — | — | — | 8 |
| Active experts/token ($k$) | — | — | — | — | 2 (top-2) |
| Expert intermediate size | — | — | — | — | 576 |
| Shared expert | — | — | — | — | Yes (sigmoid gate) |
| Shared expert hidden state | — | — | — | — | 576 |
| Load balancing | — | — | — | — | Bias + aux loss |
| Aux loss weight | — | — | — | — | 0.1 |
| *Parameters* |  |  |  |  |  |
| Total params (inference) | 99,845,376 | 38,238,720 | 100,736,256 | 37,793,280 | 102,703,360 |
| Total params (Million) | ~99.8M | ~38.2M | ~100.7M | ~37.8M | ~102.7M |
| Total params w/ MTP | — | — | — | — | ~120.9M |
| Active params/token | ~99.8M | ~38.2M | ~100.7M | ~37.8M | ~38.8M |
| KV cache elems/layer/pos | 1,536 | 1,024 | 512 | 512 | 64 |
| KV reduction vs GPT2 peer | 1× | 1× | 3× | 2× | 24× |

#### 3.3.1 Multi-Head Attention (MHA)

Multi-Head Attention (MHA) [41] is the canonical attention mechanism and serves as our dense baseline. Each of the *H* heads maintains its own query, key, and value projections of the full model dimension *d*, computes scaled dot-product attention independently, and the per-head outputs are concatenated and linearly projected back to *d* (Figure 1(e)). We adopt GPT-2 [43] as the representative model of this class, a dense, decoder-only transformer that couples MHA with a feed-forward network (FFN) using GELU activation [44], pre-layer normalization [45], and learned absolute position embeddings. In vanilla MHA, query head maintains a dedicated key and value projection, the KV cache grows as 2 × *H* × *d*_head_ elements per token position (Table 2). During training, all intermediate query, key, and value activations are reshaped [*B, H, T, d*_head_] for each of the three projections and must be retained for backpropagation, alongside the full [*B, H, T, T* ] attention-score matrix required for the softmax backward pass. This latter term scales as O(*B* · *H* · *T* ^2^) where batch size B, sequence length T,O is big-O notation, making it the dominant memory cost at large sequence lengths. At inference, gradients are absent and attention is computed autoregressively, only the accumulated KV cache, growing as 2 × *L* × *H* × *t* × *d*_head_ per sequence where *t* ∈ {1*, …, T* }, is held in memory, reducing the footprint to O(*L* · *H* · *T* ).

#### 3.3.2 Grouped-Query Attention (GQA)

Grouped-Query Attention (GQA) [29] is an attention mechanism that interpolates between MHA and multi-query attention to reduce the key-value footprint. Whereas standard MHA maintains *H* independent key and value projections, one per query head, GQA reduces this to *G* shared KV groups, where *G < H*. Each group serves *H/G* query heads (Figure 1(f)), so attention capacity is largely preserved while the number of key and value parameters shrinks by a factor of *H/G*. We adopt LLaMA-2 [39] as the representative model of this class, which pairs GQA with a set of parameter-efficient components in place of the GPT-2 counterparts. LayerNorm is replaced by RMSNorm [46], which skips mean-centring and is cheaper to compute. Learned absolute position embeddings are replaced by Rotary Position Embeddings (RoPE) [47], which encode token position by rotating the query and key vectors inside each attention head rather than adding a fixed embedding to the input. The GELU feed-forward block is replaced by SwiGLU [48], which introduces a learned gate that controls how much of the intermediate representation is passed forward:

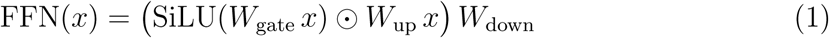

where *W*_gate_, *W*_up_, and *W*_down_ are learned weight matrices, SiLU is the sigmoid-gated linear unit, and ⊙ denotes elementwise multiplication. At the 100M tier we use *G* = 4 and *H* = 12 (a 3× reduction), which requires two additional layers (*L* = 16 versus *L* = 14 for GPT-2 at 100M) to recover the reduced parameter count and match the total budget.

The memory consequence of GQA operates differently at training and inference. During training, the full attention-score matrix [*B, H, T, T* ] is still computed and retained across all *H* query heads for the softmax backward pass, so training memory scales identically to GPT-2 at O(*B* · *H* · *T* ^2^). The benefit is realised entirely at inference, where only *G* key and value tensors are cached per layer instead of *H*. The KV cache therefore grows as 2 × *G* × *d*_head_ elements per token position (Table 2), yielding a 3× reduction at the 100M tier and a 2× reduction at the 38M tier relative to the GPT-2 peer at matched parameter budget.

#### 3.3.3 Multi-Head Latent Attention with Mixture-of-Experts (MoE-Bind)

This architecture combines two mechanisms, Multi-Head Latent Attention (MLA), which shrinks the key-value cache by compressing the attention representation rather than dropping heads, and a sparse Mixture-of-Experts (MoE) feed-forward layer, which decouples total from active parameter count. A transformer block containing the two with RMSNorm and residual skip connections (Figure 1(b)). We adopt DeepSeekV3 [24] as the representative model of this class, using MLA in place of MHA or GQA, an MoE layer in place of a dense FFN and RMSNorm.

**Multi-Head Latent Attention.** MHA and GQA reduce KV memory by operating over fewer heads, MHA caches *H* full-rank KV projections per token position, GQA caches *G < H*. MLA instead compresses the representation itself (Figure 1(c)). A learned down-projection *W^DKV^* maps each hidden state *x* ∈ R*^d^* (model dimension d) to a single low-rank key–value latent *C^KV^* ∈ R*^rKV^* (*r_KV_* = 64); paired up-projections *W^UK^* and *W^UV^* later decompress this latent back into full-rank key and value tensors across all *H* heads(*W^D^* and *W^U^* correspond to down and up projections):

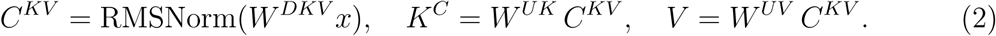

Intuitively, the down-projection performs the compression and the up-projections the reconstruction, so only the compact latent, not the full per-head key and value tensors, needs to be retained. At inference, only *C^KV^* (64 elements per token position) is cached per layer, reducing the KV cache to *r_KV_* = 64 elements per token position, a 24× reduction relative to GPT2-100M and 8× relative to LLaMA2-100M (Table 2).

Queries follow the same path through a separate down-projection into their own latent *C^Q^* ∈ R*^rQ^*(*r_Q_* = 64) before an up-projection restores them (Figure 1(c)); compressing the query lowers activation memory during training. The attention-score matrix [*B, H, T, T* ] is still retained in full for the backward pass, so training memory scales as O(*B* · *H* · *T* ^2^), the same as GPT-2 and LLaMA2 at matched head count.

**Decoupled Rotary Position Embeddings.** RoPE cannot be applied directly to the compressed latent, rotating *C^KV^* would entangle positional information with the content representation and prevent correct decompression. MLA therefore decouples position from content. Separate RoPE projections produce dedicated *r*_rope_ = 32-dimensional positional subvectors *Q*_rope_ and *K*_rope_ per head, to which RoPE is applied exclusively, while the content latent carries no position. The final query and key are formed by concatenating the content and positional parts (Figure 1(c)):

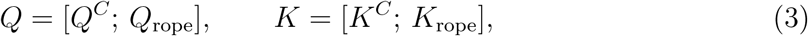

yielding an effective attention dimension of 96 per head (*d*_head_ +*r*_rope_ = 64 +32) rather than the 64 used by GPT-2 and LLaMA2. This separation is precisely what lets MLA keep its cache small while still encoding position, only the content latent *C^KV^* is cached, and the small positional key *K*_rope_ is projected directly from *x* at each step rather than stored. Unlike LLaMA2, where RoPE is applied to the full key vector, *K*_rope_ here is not part of the cached latent.

**Mixture-of-Experts Feed-Forward Layer.** The dense FFN is replaced by a sparse MoE layer consisting of *N* = 8 independent SwiGLU expert networks (Figure 1(d)). A linear router maps each token representation to *N* logits; top-*k* selection (*k* = 2) identifies the two highest-scoring experts, and the final output is the softmax-weighted sum of their outputs:

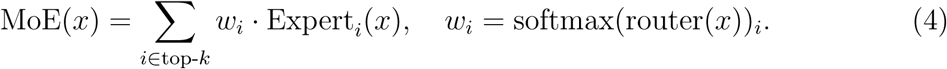

Because each token is processed by only 2 of the 8 experts, per-token compute stays low while total model capacity grows with the number of experts. An additional shared expert, applied unconditionally to every token with a learned sigmoid gate (Figure 1(d)), provides a stable residual pathway independent of routing decisions. Load balancing is enforced through a dual mechanism: a differentiable auxiliary loss L_aux_ = *N _i_ f_i_P_i_* (weight *α* = 0.1, where *f_i_* is the fraction of tokens routed to expert *i* and *P_i_* the mean routing probability) encourages uniform expert utilisation; expert-specific bias terms are updated non-differentiably at each step to correct persistent routing imbalance.

The MoE structure decouples total from active parameter count. Of the ∼102.7M total parameters, only ∼38.8M are active per forward token, the always-active components (MLA projections, norms, embeddings, router, shared expert) plus the *k/N* = 2*/*8 fraction of routed expert parameters selected per token. This is the active parameter budget used in comparative analysis against the dense 38M-tier models.

### 3.4 Pre-training Strategy

All models are pre-trained on UniRef50 [30] using a next-token prediction objective, minimising cross-entropy loss over every token in the input sequence. Training configuration details are given in Table 3.

**Table 3:** Complete pre-training hyperparameters for all five models.

|  | MHA-100M | MHA-38M | GQA-100M | GQA-38M | MoE-Bind-100M |
| --- | --- | --- | --- | --- | --- |
| **Hyperparameters** |  |  |  |  |  |
| Token budget | 200M | 76M | 200M | 76M | 200M |
| Micro-batch size | 28 | 52 | 6 | 12 | 8 |
| Gradient accum steps | 18 | 5 | 85 | 21 | 64 |
| Effective batch (seqs) | 504 | 260 | 510 | 252 | 512 |
| Eff. batch (tokens/update) | 387,072 | 199,680 | 391,680 | 193,536 | 393,216 |
| Optimizer updates | 517 | 381 | 510 | 393 | 510 |
| Warmup steps | 186 | 38 | 868 | 164 | 650 |

Protein sequences are tokenised at the character level, assigning one token per amino acid residue (vocabulary size 31: 20 standard amino acids plus special tokens). Subword tokenization methods such as BPE [49], standard in natural language processing, are not appropriate for protein sequences, amino acids are already the irreducible biochemical units of a protein, and merging residue pairs into subword tokens would destroy the one-token-one-residue correspondence without any meaningful compression benefit given the small natural vocabulary size.

Each model is trained to 10% of its Chinchilla-optimal token budget [50] because of resource constrains, 200M tokens for the 100M-parameter tier and 76M tokens for the 38M tier. This is a deliberate training choice, the objective is architectural comparison under matched compute constraints, not maximising absolute perplexity. for all the models, learning rate is 10^−4^, and minimul learning rate is 10^−5^, learning-rate schedular is cosine decay. The same random seed is used across all models to control for stochasticity in training dynamics and ensure that observed differences are due to architecture rather than random variation.

Multi-Token Prediction (MTP) [51] augments the standard next-token objective with auxiliary prediction heads that forecast *k* future tokens simultaneously during pre-training. To assess whether MTP improves protein sequence pre-training, MoE-Bind is pre-trained under two conditions: MTP-ON (2 auxiliary heads, loss weight 0.3) and MTP-OFF with standard next token prediction objective. MTP heads are discarded before fine-tuning in both cases. MHA and GQA are trained without MTP.

### 3.5 Fine-tuning for Binder Generation

Pre-trained language models acquire a general prior over protein sequence statistics, amino acid frequencies, sequence characteristics and patterns, but are not optimized for any specific generative task. Fine-tuning adapts these pre-trained weights to the conditional binder generation objective by continuing training on experimentally validated protein–protein interaction pairs, where the model must learn to generate a plausible binder sequence given a target sequence as context. Without this task-specific stage, a pre-trained model can only perform unconditional sequence generation and has no mechanism to condition its output on a particular target. Fine-tuning introduces this conditioning by exposing the model to paired (*X, Y* ) examples in which the target sequence *X* precedes the binder *Y* , so that autoregressive generation of *Y* is conditioned on *X*, shifting the learned distribution from *p*(*y_t_* | *y*_1:_*_t_*_−1_) toward *p*(*y_t_* | *y*_1:_*_t_*_−1_*, X*).

Protein sequences are tokenized at the character level, assigning one token per amino acid residue. The vocabulary contains 31 tokens: 20 standard amino acids, 5 rare or ambiguous residue codes (X, B, Z, U, O), two control tokens (<PAD>, <EOS>), and four structural delimiter tokens (<PROTEIN_A>, </PROTEIN_A>, <PROTEIN_B>, </PROTEIN_B>). Models are fine-tuned on PPI sequence pairs using the conditional sequence generation objective defined in Section 3.1. The input to the model is the concatenated sequence:

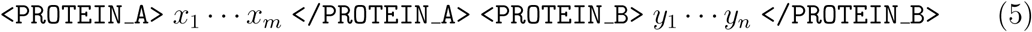

where the four structural delimiter tokens explicitly encode the role of each subsequence. By default, the cross-entropy loss is computed over the entire concatenated sequence, gradients are taken from both the target tokens *X* (within <PROTEIN_A> . . . </PROTEIN_A>) and the binder tokens *Y* (within <PROTEIN_B> . . . </PROTEIN_B>). The model therefore learns to model the joint target-binder sequence, while the structural delimiters and the left-to-right ordering supply the target-conditioning signal at generation time. We refer to this default setting as conditional masking disabled (CM-False), and it is the framework used throughout unless stated otherwise. At inference, the model receives the target prefix up to and including <PROTEIN_B> and generates the binder sequence autoregressively until </PROTEIN_B> or <EOS> is produced.

As an ablation we additionally consider conditional masking enabled (CM-True), in which the loss is restricted to the binder tokens *Y* and all target tokens are masked from the loss, so that gradient updates derive only from binder prediction. The effect of this choice-together with the MTP ablation-on binder quality is examined in the peptide binder generation analysis (Section 4.4). Fine-tuning is run for a fixed sample budget across all architectures (Table 4), so any observed differences in performance are attributable to architecture and initialisation rather than optimisation configuration. The learning rate is set an order of magnitude below the pre-training rate to prevent catastrophic forgetting of protein sequence priors acquired during pre-training. For the pre-train + fine-tune regime, fine-tuning is initialised from the pre-trained checkpoint; for the Only-FT regime, weights are initialised randomly with the same configuration. MTP auxiliary heads in MoE-Bind are disabled before fine-tuning in both cases and contribute no gradient updates during this stage.

**Table 4:**
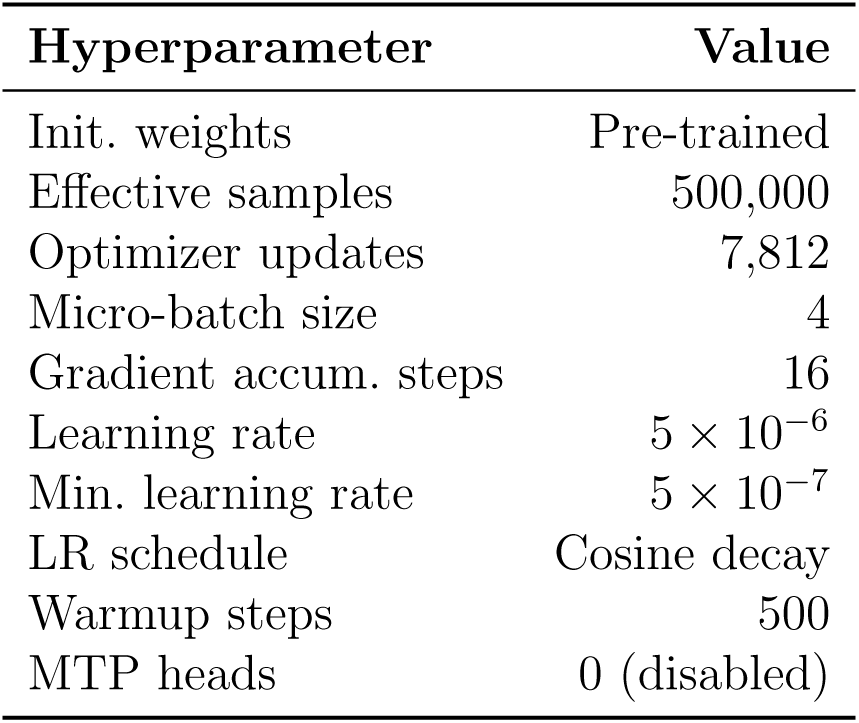
Fine-tuning hyperparameters, identical across all five models

| Hyperparameter | Value |
| --- | --- |
| Init. weights | Pre-trained |
| Effective samples | 500,000 |
| Optimizer updates | 7,812 |
| Micro-batch size | 4 |
| Gradient accum. steps | 16 |
| Learning rate | $5 \times 10^{-6}$ |
| Min. learning rate | $5 \times 10^{-7}$ |
| LR schedule | Cosine decay |
| Warmup steps | 500 |
| MTP heads | 0 (disabled) |

### 3.6 Computational Infrastructure

All models were implemented in PyTorch with CUDA 12.1. Model development, inference, and the routing and sequence-level analyses were carried out on a local workstation with a single NVIDIA RTX 3060 GPU (12 GB). Some portion of training were performed on an NVIDIA L40 GPU (48 GB) provided through the India AI Compute initiative (Project ID: P1-S2025070964).

## 4 Results

### 4.1 Sequence level evaluation

We evaluated the compositional and physicochemical properties of binders generated for the 22-sequence DB5 benchmark using the 100M-parameter variants of MHA, GQA, and MoE-Bind. This DB5 benchmark was constructed to be leakage-free with respect to both the pre-training and fine-tuning datasets. DB5 proteins that were similar to sequences in either the UniRef50 pre-training corpus or the STRING fine-tuning corpus were removed using MMseqs2 at a 10% sequence-identity cutoff and 80% bidirectional coverage, leaving 22 leakage-free DB5 pairs for evaluation. We compared the generated binders against two reference distributions, the DB5 ground-truth binders and a 1,000-sequence random sample from the STRING fine-tuning corpus. Figure 2 summarizes four complementary sequence-level properties.

**Figure 2:**
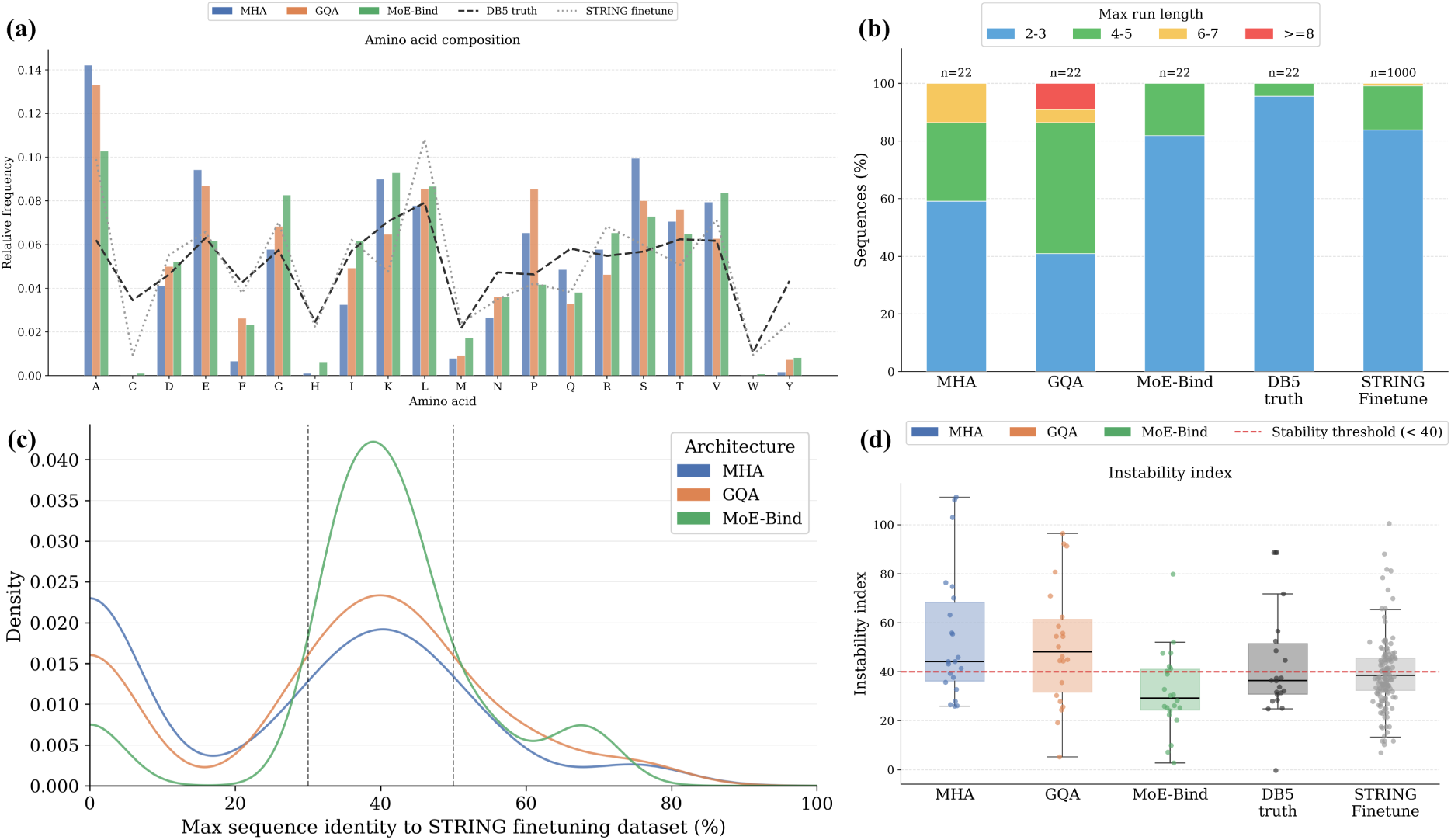
Sequence-level characterization of binders generated by the 100M-parameter MHA, GQA, and MoE-Bind models on the leakage-free DB5 benchmark. The generated sequences are compared against the DB5 ground-truth binders and a 1,000-sequence random sample from the STRING fine-tuning corpus. **(a)** Amino acid composition of generated binders relative to DB5 and STRING reference distributions. **(b)** Maximum consecutive single-amino-acid run-length composition, grouped into 2–3, 4–5, 6–7, and≥ 8 residue runs. **(c)** Sequence novelty measured as the maximum sequence identity between each generated binder and the STRING fine-tuning corpus; dashed vertical lines indicate the intermediate novelty window. **(d)** Instability-index distributions, where the red dashed line marks the stability threshold of 40; sequences below this threshold are predicted to be stable.

**Amino acid composition** All three architectures(Figure 2a) produce binder sequences whose amino acid composition closely relate the DB5 ground-truth amino acids, where the dataset entirely held out from both pre-training and fine-tuning, reflecting genuine generalization rather than memorization. The models were fine-tuned exclusively on the much larger STRING PPI corpus, yet their generated sequences align with the DB5 reference comparably well, indicating that the compositional statistics learnt from STRING transfer faithfully to unseen natural interfaces. MHA and GQA both correctly captured the presence of hydrophobic, polar, and charged residues that define natural binders. MoE-Bind’s gets even more closely to the DB5 ground truth across the full amino acid alphabet, suggesting that its MoE-Bind architecture extracts a richer and more transferable representation of interface composition from the STRING fine-tuning data. MoE-Bind additionally shows a mild enrichment of basic residues (Arg, Lys) relative to the DB5 reference, a subtle compositional signature that is entirely consistent with the electrostatic character of natural protein interfaces.

**Homopolymer run-length composition** Figure 2b shows the maximum consecutive run length in each sequence. Here, run length indicates the longest stretch of the same amino acid repeated without interruption, such as “AAAA” having a run length of 4.

Long single-residue repeats correspond to homopolymeric or low-complexity segments, which are commonly treated as a warning sign in protein sequence analysis because they can reflect compositionally biased sequence generation rather than natural protein-like design [52, 53]. This is especially important for autoregressive protein language models, where repetitive outputs are a known generation failure mode and repetition penalties are often used to reduce such behavior [17]. Although some natural low-complexity regions can be functional, excessive homopolymer runs are generally not ideal for protein sequence generation because they may reduce globular protein-like character and biological plausibility.

The DB5 ground-truth binders show the cleanest pattern, with nearly all sequences (∼95%) having short runs of only 2–3 residues and only a small fraction reaching 4–5 residues. The STRING fine-tuning set shows a similar natural pattern, most sequences (∼84%) also fall in the 2–3 range, some have 4–5 residue runs, and only a very small fraction reach 6–7 residues. Importantly, neither DB5 nor STRING contains sequences with very long runs of ≥ 8 residues.

MHA produces mostly short runs, with nearly 60% of sequences in the 2–3 range, but it also has a noticeable fraction with longer runs, about 27% fall in the 4–5 range and about 14% reach 6–7 residues. This suggests that MHA is generally controlled, but it sometimes generates more repetitive regions than expected from true binder sequences. GQA shows the least ideal pattern among the three models. Only about 41% of its sequences remain in the short 2–3 range, while nearly half fall in the 4–5 range, and around 9% contain very long runs of ≥ 8 residues. This indicates a stronger tendency toward repetitive or low-complexity outputs. MoE-Bind gives the best model pattern, with about 82% of sequences in the 2–3 range and the remaining ∼18% only in the 4–5 range. It does not generate any 6–7 or ≥ 8 residue runs, making its run-length profile closest to the natural DB5 and STRING distributions.

**Sequence novelty relative to fine-tuning data** Sequence novelty plot(Figure 2c) show how similar the generated binders are to the STRING fine-tuning sequences, measured by maximum sequence identity. All three models generate sequences that are clearly different from the fine-tuning dataset, suggesting that they are not merely copying training examples. MHA and GQA have broader distributions with one peak near 40% identity and another similar peak show higher density near 0% identity, meaning that some of their generated sequences are extremely distant from any sequence in the STRING fine-tuning set. While this indicates high novelty, it may not be ideal because sequences that are too far from known protein sequence space can be less likely to preserve realistic protein-like patterns and characteristics. In contrast, MoE-Bind shows a narrower distribution, mostly centered around 35–45% identity and lowest density at 0% identity. This means that MoE-Bind generates sequences that are still novel, but they remain closer to the sequence space learned from STRING. Overall, MoE-Bind appears to produce more consistently controlled outputs, while MHA and GQA generate a wider range of sequence novelty, including more very low-identity sequences.

**Physicochemical stability** Figure (Figure 2d) shows the instability index of the generated and reference sequences. The instability index is a sequence-based estimate of protein stability calculated from dipeptide composition. Following the original definition by Guruprasad et al., proteins with an instability index below 40 are predicted to be stable, whereas values above 40 suggest possible instability [54]. This metric is applicable here because generated binders should not only be novel, but should also remain compatible with stable protein-like behavior.

The DB5 ground-truth binders provide the natural reference for true binding sequences. Their median instability index is below the stability threshold, around 36, and approximately two-thirds of the sequences fall below 40. This shows that most DB5 binders are predicted to be stable. The STRING fine-tuning set shows a similar but slightly broader pattern, with a median instability index close to the threshold, around 38–39, and a little over half of the sequences below 40. This indicates that the fine-tuning data contains many stable sequences, but also includes a wider range of stability values. MHA has a median instability index above the stability threshold, around 44, with only about one-third of its sequences falling below 40. This means that a large fraction of GPT2-generated binders are predicted to be less stable than the DB5 and STRING references. GQA shows the weakest stability profile among the three models, with the highest median instability index, around 48, and again only about one-third of sequences below the stable threshold. This suggests that GQA produces more sequences that may be unstable. MoE-Bind gives the best stability profile, with the lowest median instability index, around 29 to 30, and roughly two-thirds of its sequences below 40. Thus, MoE-Bind produces more sequences predicted to be stable than MHA and GQA, and its stability profile is closest to, or slightly better than, the DB5 reference.

### 4.2 Structure-level binder interface confidence

We evaluated whether the generated binders form plausible complexes with their true receptors at the structure level. The primary evaluation was performed on the 22 leakage-free DB5 benchmark pairs, which are filtered for sequence overlap against both the pre-training and fine-tuning corpora. For each target, the receptor sequence was used as the conditioning input sequence, and each model generated a binder with the same length as the native DB5 ligand. This setup tests whether a model can generate a binder for the same receptor, rather than simply reconstructing the original DB5 binding partner. The binder generation process included a quality-filtering step to remove low-quality candidates, for each target, 10 binders were sampled and then filtered based on maximum single-residue run length, sequence repetitiveness, and model perplexity. The single best surviving candidate was paired with the corresponding DB5 receptor and evaluated structurally.

For each ligand-receptor pair, we predicted the complex structure using ColabFold with AlphaFold2-Multimer v3. We used the interface predicted TM-score (ipTM) as the main structural confidence metric because it estimates the confidence of the predicted interaction between chains. For each DB5 target, the native receptor-ligand pair was also folded using the same ColabFold protocol to obtain a reference ipTM, and each generated complex was compared directly against this reference.

To test whether the trends observed on the 22-target set are robust to benchmark size and to the choice of structure predictor, we repeated the evaluation on a larger benchmark of 121 DB5 pairs filtered for leakage against the fine-tuning data (STRING physical interactions, 10% sequence identity). For each of these targets every model generated a receptor-conditioned binder using the same sampling and quality-filtering procedure, and both the 121 generated complexes per model and the 121 native reference complexes were folded with Boltz-2 using multiple sequence alignments (MSA). We selected Boltz-2 for this larger run because it provides co-folding accuracy approaching that of AlphaFold3 when supplied with MSA input, while being substantially faster and more practical for the volume of predictions required here (more than 700 complexes), and because it is an architecture and training pipeline entirely distinct from AF2-Multimer, providing an independent structural cross-check rather than a correlated re-run of the same model.

A subset of complexes exceeded the available GPU memory under Boltz-2 with MSA and could only be folded with a no-MSA fallback. Because such fallback predictions are produced by a different protocol, are not on the same confidence scale, and occur unevenly across models, mixing them with MSA-based scores would confound the comparison. We therefore retained only those targets for which the native reference and all five models produced a successful Boltz-2 MSA prediction. The native references folded under Boltz-2 MSA for 84 of the 121 targets, and the intersection with all five models leaves a shared, fully paired set of 82 targets scored under a single uniform protocol. Within this set we further found three groups of targets that, although listed under different identifiers, comprised identical receptor and binder sequences (i.e. the same biological complex submitted more than once). Retaining all copies would both multiply-count these complexes and let a single stochastic reference fold set the hit threshold, so we collapsed each group to one entry: the pass threshold was taken as the mean reference ipTM over the group’s folds, and each model’s representative was its lowest-perplexity generated binder for that complex. This deduplication yields the final set of 78 unique targets, on which all Boltz-2 results are reported.

A generated binder was counted as a structural hit when its predicted ipTM was greater than or equal to the reference DB5 complex ipTM:

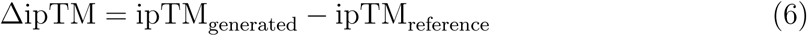

Thus, a hit corresponds to ΔipTM ≥ 0. This is a strict criterion because the generated binder must reach or exceed the structural confidence of the native DB5 receptor-ligand complex for the same target.

In this structure-level experiment, we compared three 100M-parameter sequence models with different attention and routing designs. MHA, a standard multi-head attention transformer; GQA, a grouped-query attention transformer; and our MoE-Bind model, which combines multi-head latent attention with sparse mixture-of-experts routing. These three models use only receptor sequence conditioning to generate the binder sequence, allowing us to test whether the MoE-Bind design improves target-conditioned binder generation over dense transformer baselines. We also included two external baselines to place the results in context. PepMLM-650M is a target-sequence conditioned peptide binder generation model based on ESM-2 and masked language modeling. This comparison is not fully balanced because PepMLM was designed and trained with binder lengths limited to 50 amino acids and target lengths limited to 500 amino acids in its curated training setup [34]. Our DB5 binders are often longer protein chains rather than short peptides. Nevertheless, PepMLM is included because it is one of the few open-sourced sequence-conditioned binder generation models, and no directly equivalent full-length, sequence-only protein binder generator model is openly available. As a structure-based baseline, we included RFdiffusion followed by ProteinMPNN. RFdiffusion is a state of the art protein binder backbone generation method, but unlike our sequence-only models it requires the receptor structure as input. In addition, RFdiffusion alone does not directly provide the final amino acid binder sequence; the generated backbone must be sequence-designed using ProteinMPNN before folding and evaluation. Therefore, the RFdiffusion with ProteinMPNN pipeline provides a useful structure-level binder-design reference point.

Figure 3 shows the generated ipTM against the reference ipTM for MHA-100M, GQA-100M, MoE-Bind-100M, PepMLM-650M, and RFdiffusion with ProteinMPNN on the 22 leakage-free DB5 targets. Each point represents one DB5 target; points above the diagonal are counted as hits and points below as misses. Targets are grouped by reference difficulty: easy targets have reference ipTM below 0.4, medium targets fall between 0.4 and 0.7, and hard targets have reference ipTM above 0.7. Most successful cases occur in the low-reference-ipTM regime, while medium and hard targets remain difficult for all evaluated methods. Within this benchmark, MoE-Bind shows the strongest structure-level performance among the tested methods, it achieves 6 hits out of 22 targets, a 27.3% hit rate, and the least negative mean ΔipTM (−0.255). MHA reaches 3 of 22 hits (mean ΔipTM −0.278) and GQA reaches 4 of 22 (mean ΔipTM −0.269), while the external baselines PepMLM-650M and RFdiffusion with ProteinMPNN each reach 4 of 22 hits. Thus, although the absolute hit rates remain modest, MoE-Bind gives the best result under the AF2-Multimer ipTM-hit criterion, with the highest number of above-diagonal points indicating more frequent recovery of complexes whose predicted interface confidence matches or exceeds the corresponding DB5 reference.

**Figure 3:**
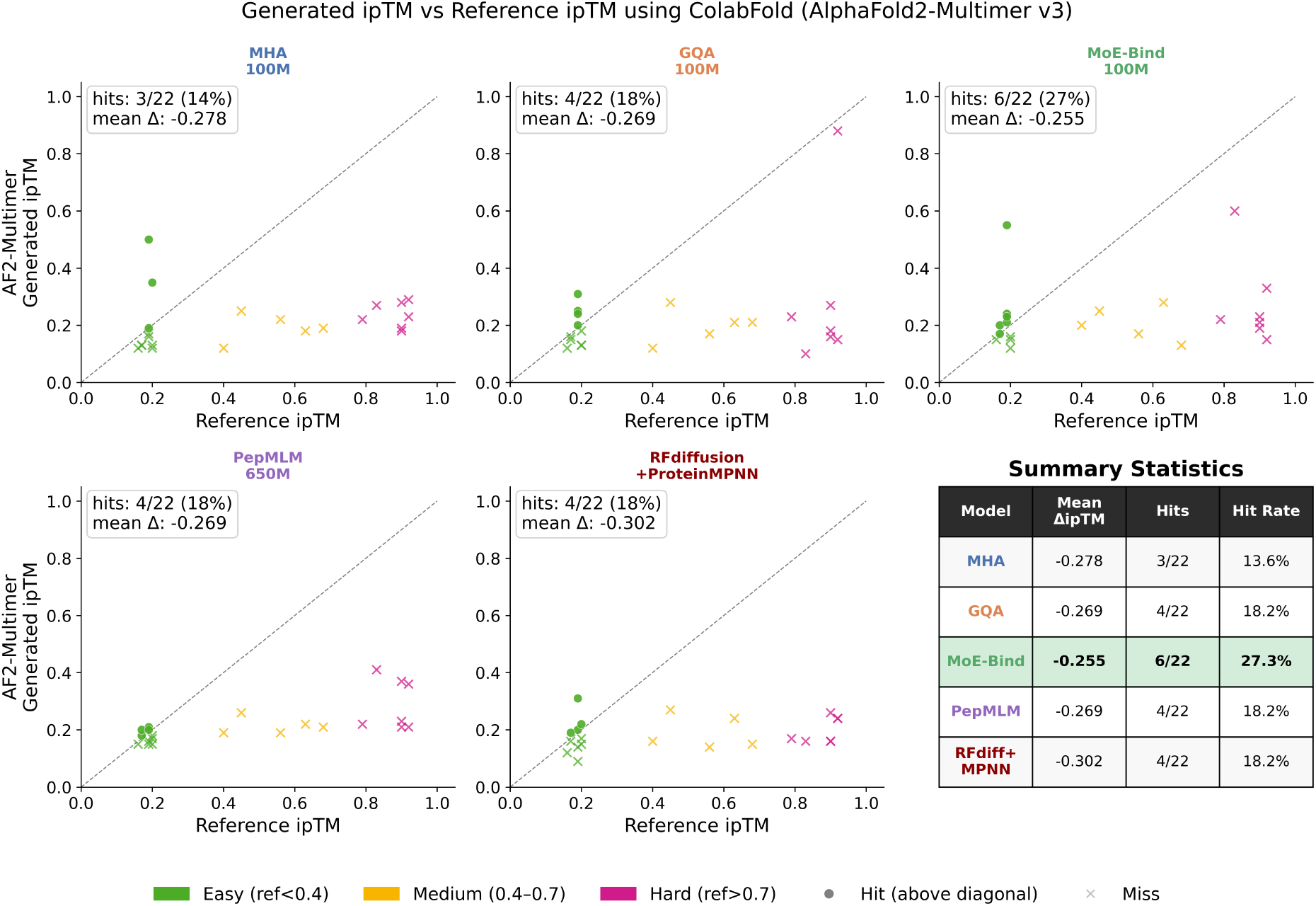
Structure-level evaluation of generated binders using ColabFold with AlphaFold2-Multimer v3. For each of the 22 leakage-free DB5 targets, the generated receptor-binder complex was compared against the corresponding DB5 reference complex using ipTM. Each scatter point represents one DB5 target, with the x-axis showing the reference-complex ipTM and the y-axis showing the generated complex ipTM. The dashed diagonal indicates the hit threshold, where generated ipTM is equal to reference ipTM; points above this line are counted as hits. Targets are colored by reference difficulty; easy, reference ipTM *<* 0.4; medium, 0.4–0.7; and hard, reference ipTM *>* 0.7. The summary table reports mean ΔipTM, number of hits, and hit rate for each method. MoE-Bind achieves the highest hit rate and the least negative mean ΔipTM among the evaluated methods in this benchmark.

This ranking is reproduced on the larger, independently folded benchmark. Table 5 reports the Boltz-2 MSA evaluation on the 78 shared targets, now including the compute-matched 38M-parameter dense models. MoE-Bind again attains the highest hit rate (24.36%), exceeding both ∼100M dense models (GQA-100M 23.08%, MHA-100M 21.79%) and both compute-matched ∼38M dense models (GQA-38M 20.51%, MHA-38M 16.67%), while activating only ∼38.8M parameters per token. The same ordering therefore holds under an independent folding engine and on a larger target set. Under the compute-fair framework MoE-Bind outperforms its dense compute-matched peers, and under the parameter-equivalent framework it matches dense models of up to 2.6× its active size at a fraction of the per-token compute. These results are therefore best read as MoE-Bind consistently matching the dense baselines across two structure predictors while operating at substantially lower active compute, rather than significantly surpassing them.

**Table 5:**
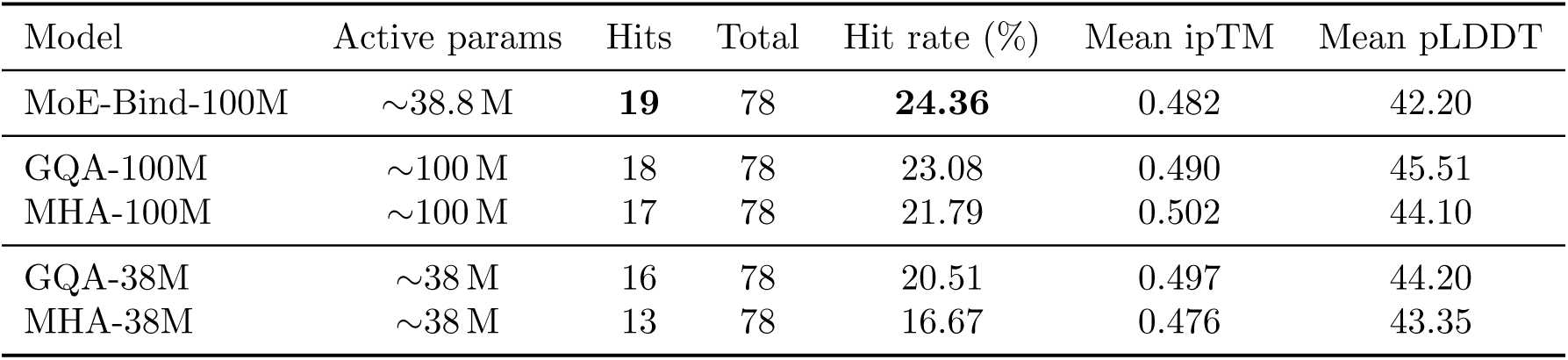
Structure-level binder evaluation under Boltz-2 with MSA on the shared subset of complexes for which the reference and all five models produced a Boltz-2 MSA prediction. Complexes that appear under multiple IDs with identical receptor and binder sequences were deduplicated to a single entry, yielding 78 unique complexes. A hit is counted when the generated complex ipTM meets or exceeds the native DB5 reference ipTM. MoE-Bind attains the highest hit rate while activating only ∼39 M parameters per token; all models produce binders with low monomer-fold confidence (pLDDT *<* 50).

| Model | Active params | Hits | Total | Hit rate (%) | Mean ipTM | Mean pLDDT |
| --- | --- | --- | --- | --- | --- | --- |
| MoE-Bind-100M | $\sim 38.8$ M | **19** | 78 | **24.36** | 0.482 | 42.20 |
| GQA-100M | $\sim 100$ M | 18 | 78 | 23.08 | 0.490 | 45.51 |
| MHA-100M | $\sim 100$ M | 17 | 78 | 21.79 | 0.502 | 44.10 |
| GQA-38M | $\sim 38$ M | 16 | 78 | 20.51 | 0.497 | 44.20 |
| MHA-38M | $\sim 38$ M | 13 | 78 | 16.67 | 0.476 | 43.35 |

### 4.3 Structural Characterization of MoE-Bind Binder Hits

Beyond the aggregate hit rate (Section 4.2), we examined individual complexes to characterize how the generated binders engage their targets. Of the 19 MoE-Bind binders that meet the Boltz-2 hit criterion (ΔipTM ≥ 0), we selected four representative cases (1EFN, 2OOB, 3H2V, 1KTZ) spanning a range of reference difficulty for visualization (Figure 4).

**Figure 4:**
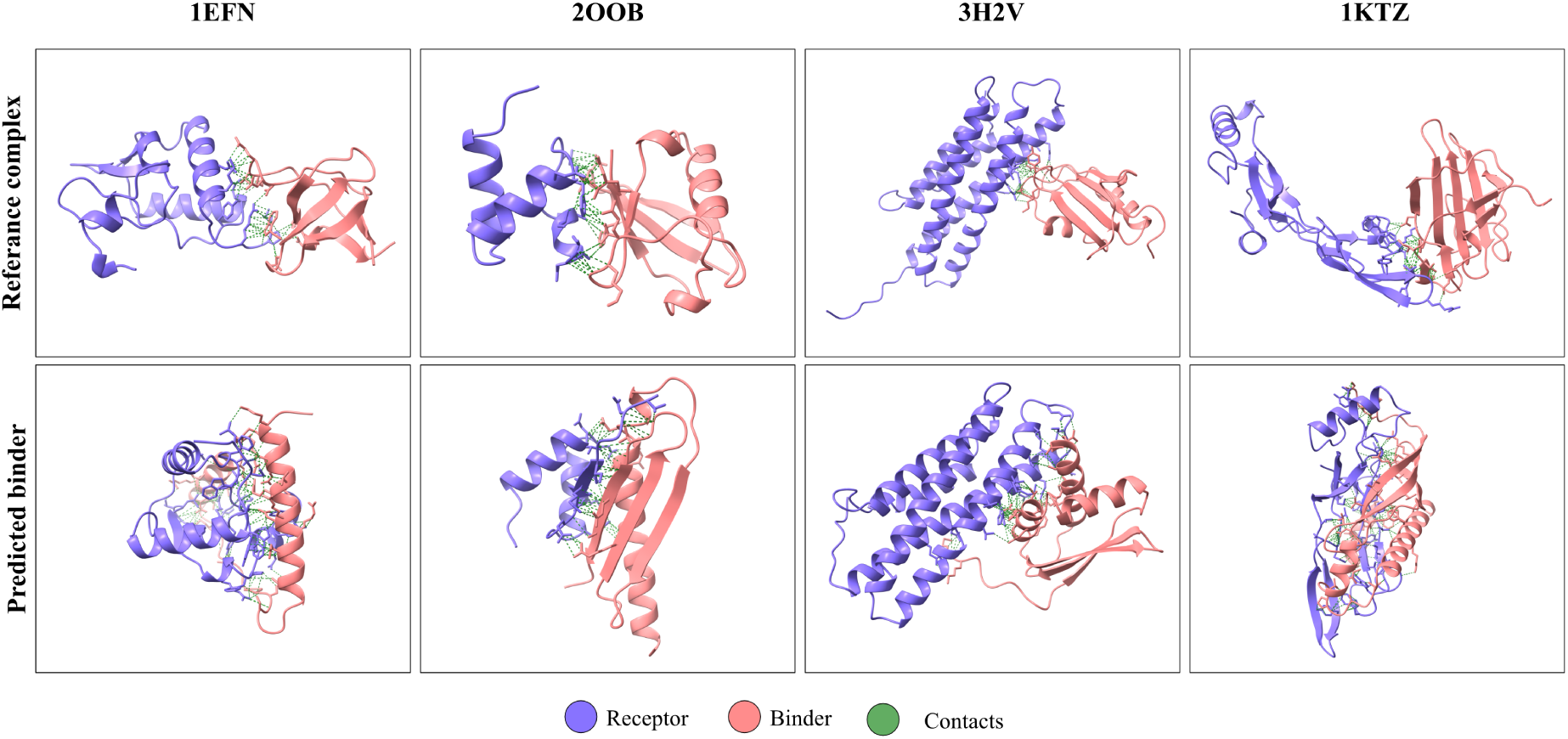
Structural characterization of four representative MoE-Bind binder hits (Boltz-2 with MSA). Columns correspond to targets (PDB IDs); the top row shows the Boltz-2 prediction of the native receptor-ligand pair (reference) and the bottom row the Boltz-2 prediction of the receptor with the MoE-Bind-generated binder. Receptor in blue, binder in red; green spheres mark inter-chain interface contacts. Both rows are Boltz-2 predictions under identical settings.

For each target, the native receptor-ligand pair and the receptor-generated-binder pair were folded under the identical Boltz-2 protocol; the native prediction provides the reference against which the ipTM hit threshold is defined. Both complexes are therefore Boltz-2 predictions, not experimental structures, and are directly comparable model outputs.In every case the generated binder adopts compact secondary structure and shows binding against the target surface, forming an interface (green contacts, Figure 4) whose predicted confidence matches or exceeds the native reference (Table 6; e.g., 1EFN ipTM 0.92 vs. 0.57).Because the generated sequences share no homology with the native binders and they produce the receptor conditioned binders de novo, which are not expected to reproduce the native epitope or fold; the comparison demonstrates plausible target engagement at comparable or higher interface confidence, not recapitulation of the natural interaction.

**Table 6:** Structure-level metrics for four MoE-Bind binder hits (Boltz-2, MSA). ΔipTM ≥ 0.

| Target | Generated | | | | Reference (native) | | | | $\Delta\text{ipTM}$ |
| --- | --- | --- | --- | --- | --- | --- | --- | --- | --- |
| | ipTM | pLDDT | BSA ($\text{\AA}^2$) | H-bonds | ipTM | pLDDT | BSA ($\text{\AA}^2$) | H-bonds | |
| 1EFN | 0.924 | 70.5 | 1586 | 7 | 0.574 | 86.2 | 481 | 6 | +0.350 |
| 2OOB | 0.852 | 57.5 | 848 | 8 | 0.755 | 94.3 | 500 | 1 | +0.096 |
| 3H2V | 0.754 | 54.6 | 1063 | 6 | 0.663 | 84.8 | 469 | 2 | +0.091 |
| 1KTZ | 0.583 | 40.9 | 2015 | 10 | 0.521 | 88.9 | 479 | 2 | +0.062 |

Quantitatively, the generated complexes form more extensive interfaces than the native references by both buried surface area (848-2015 Å^2^ vs. 469-500 Å^2^) and inter-chain hydrogen bonds (6-10 vs. 1-6). Since the native references are themselves de novo Boltz-2 predictions that form comparatively small interfaces (≈480 Å^2^, 1-6 hydrogen bonds); the comparison therefore reflects same-protocol predictions rather than experimental native interfaces. Buried area also scales with binder size and chain extension, and the largest interface here 1KTZ at 2015 Å^2^ corresponds to the lowest-confidence binder (pLDDT 40.9), where partial disorder likely inflates the contact area. We accordingly read buried-area and hydrogen-bond counts as qualitative descriptors of interface extent on predicted structures rather than as quantitative affinity estimates. Consistent with the sequence-only training objective, generated binders show modest monomer-fold confidence (pLDDT ≈41-71). We note that this is a property of the generation paradigm rather than the MoE active-parameter budget, all architectures in this study, including the dense 100M baselines, produce binders in a similar pLDDT range (Section 4.2). Despite this modest fold confidence, MoE-Bind attains interface confidence (ipTM) competitive with the dense baselines while activating only ∼38.8M parameters per token.

### 4.4 Peptide-binder interface comparison

In addition to full-length protein binder generation, we evaluated whether the sequence-conditioned generative models can transfer to short peptide binder design. Peptides are short amino-acid chains, usually shorter than protein binders, and are attractive because they can bind protein surfaces that are difficult to target with small molecules. This setting is deliberately out-of-distribution for our models, MoE-Bind, MHA, and GQA were trained only from sequence data for general-length protein generation and were not specifically optimised for peptides of length ≤ 50. The comparison to PepMLM-650M is therefore not intended as a strictly fair head-to-head benchmark. PepMLM is a peptide-specific, target-sequence-conditioned masked language model, finetuned from ESM-2 to reconstruct peptide binders appended to target protein sequences [34]. Instead, this experiment tests whether the MoE-Bind architecture retains useful target-conditioned binding signal in a short-peptide regime that it was not explicitly trained for. We used the PepMLM test set and removed targets with high sequence similarity to our finetuning data using MMseqs2 filtering, leaving 68 peptide-target pairs. For each target, models generated a peptide of the same length as the reference peptide. Because this comparison spans nine models, PepMLM-650M, four MoE-Bind variants, and the dense MHA and GQA baselines, across 68 targets, all peptide-target complexes were predicted with Boltz-1 in sequence-only mode, which is substantially faster than Boltz-2 with MSA and made the large number of predictions tractable; every complex was additionally re-folded with AlphaFold2-Multimer as an independent structural cross-check. All models, including PepMLM-650M, were evaluated under identical protocols, and hits were defined relative to references folded with the same tool, so the comparison reflects relative trends rather than cross-protocol absolute scores. Because short-peptide design is out-of-distribution for our models, we did not assume that the configuration optimal for full-length binders (MTP-ON, CM-False) would remain optimal here. We therefore evaluated all four combinations of conditional masking, CM-False (training loss over the full target-binder sequence) and CM-True (loss over the binder tokens only) and multi-token prediction (MTP-ON/MTP-OFF), to probe how each training choice transfers to the short-peptide regime rather than presupposing a single default. The reference score for each target is the predicted ipTM of the original peptide-target pair. We define an ipTM hit as stated in Equation 6, where the generated peptide-target ipTM must be greater than or equal to the reference peptide-target ipTM; that is, a generated peptide is counted as a hit if its predicted interface confidence is at least as high as that of the original reference peptide for the same target. Aggregate hit rates for all architectures under the matched conditional-masking settings are reported in Table 7, and the per-target and difficulty-stratified analyses in Figure 5. Figure 5a shows generated ipTM against reference ipTM for PepMLM-650M and the MoE-Bind variants. Points above the diagonal are hits. As expected, PepMLM performs best overall, with 35/68 hits using Boltz-1 and 29/68 hits using AF2-Multimer, consistent with PepMLM being directly trained for peptide binder generation. Among the MoE-Bind models, the best overall variant is MTP-OFF CM F, which achieves 30/68 Boltz-1 hits and 23/68 AF2-Multimer hits. The remaining MoE-Bind variants are lower, with Boltz-1 hit rates between 25/68 and 27/68 and AF2-Multimer hit rates between 18/68 and 23/68. The lower absolute performance of MoE-Bind compared with PepMLM is expected because this benchmark uses short peptides, whereas our models were not peptide-specialised. Nevertheless, the MoE-Bind models recover a substantial fraction of reference-level peptide binders, indicating that the target-conditioned sequence generation objective transfers partially to peptide binding even without peptide-specific training. An important trend is that MTP-OFF performs better than MTP-ON in this peptide benchmark, the opposite of what we observe in longer binder settings. Multi-token prediction is beneficial when generation requires longer-range planning across many residues, because the model learns representations that predict multiple future tokens simultaneously. For very short peptides, however, there is little long-range sequence context to exploit, and the simpler next-token objective used by MTP-OFF appears better calibrated for short autoregressive generation, whereas the MTP objective is more useful for longer protein-scale binders. This length-dependent behaviour is the primary reason MTP-ON remains our default for full-length binder generation while MTP-OFF is the stronger configuration here. Figure 5b stratifies AF2-Multimer hit rate by target difficulty, where difficulty is defined using the reference ipTM. Easy targets have reference ipTM *<* 0.4, medium targets between 0.4 and 0.7, and hard targets *>* 0.7. PepMLM is strongest on easy and medium peptide targets, reaching 79% and 55% hit rate respectively. MoE-Bind variants are competitive on easier targets, with MTP-ON CM T also reaching 79% on the easy tier and MTP-OFF CM T reaching 64%. On hard targets all models drop substantially, PepMLM retains the highest hard-tier AF2-Multimer hit rate at 19%, while the best MoE-Bind variants reach 19% or 9% depending on configuration. Because hard targets are those with the highest-confidence native interfaces rather than the longest peptides, this drop is difficulty-driven rather than length-driven, and it indicates that high-confidence peptide interfaces remain challenging for sequence-only generation, especially for models not trained specifically on peptide binders. Figure 5c shows the same tiered analysis using Boltz-1 and additionally includes the dense MHA and GQA baselines. In this evaluator, the best MoE-Bind variant in each tier remains competitive with PepMLM and exceeds the corresponding dense MHA/GQA variants, though the winning MoE-Bind configuration changes across tiers. MoE-Bind MTP-OFF CM F has the highest easy-tier Boltz-1 hit rate at 71%, exceeding PepMLM’s 43% under the Boltz-1 evaluator. For medium targets, PepMLM reaches 50% while MoE-Bind MTP-OFF CM F reaches 45%. For hard targets PepMLM is strongest at 56%, but MoE-Bind MTP-ON CM T reaches 47%, and several MoE-Bind and dense baselines remain in the 31-38% range. Aggregated over all 68 targets (Table 7), the comparison at a matched conditional-masking setting confirms the same ordering, at CM-False, MoE-Bind (MTP-OFF) attains a 44.1% Boltz-1 hit rate, exceeding GQA (38.2%) and MHA (30.9%) at equal total parameters, while at CM-True the four configurations cluster within a narrow 36.8-39.7% band. The MoE-Bind advantage over the dense baselines is therefore clearest at CM-False and becomes statistically indistinguishable at CM-True. We emphasise that no single MoE-Bind configuration dominates across all difficulty tiers, MTP-OFF CM F is strongest on easy targets, whereas MTP-ON CM T is strongest on hard targets. The consistent observation is not the specific model level, instead are on the architecture-level, within each difficulty regime and under each evaluator, and in the aggregate hit rates of Table 7, the best MoE-Bind variant matches or exceeds the specialised PepMLM baseline in the easier regimes and surpasses the corresponding dense MHA and GQA variants, which themselves exhibit analogous configuration sensitivity. Because peptide generation lies outside the training distribution, this configuration spread is expected different training objectives transfer differently to an unseen regime and we interpret the result as evidence for the capacity of the MoE-Bind architecture family, not of any particular checkpoint. To avoid favourable selection, we report every variant for every architecture rather than only the best-performing one. Overall, the peptide benchmark should be interpreted as an out-of-distribution transfer experiment rather than a direct replacement for PepMLM. PepMLM remains the specialised peptide-generation baseline, but MoE-Bind produces a meaningful number of reference-level peptide binders without any peptide-specific training. The MTP trend further clarifies a length-dependent behaviour, MTP helps longer binder generation, while MTP-OFF is better suited to very short peptide generation where single-token calibration dominates.

**Figure 5:**
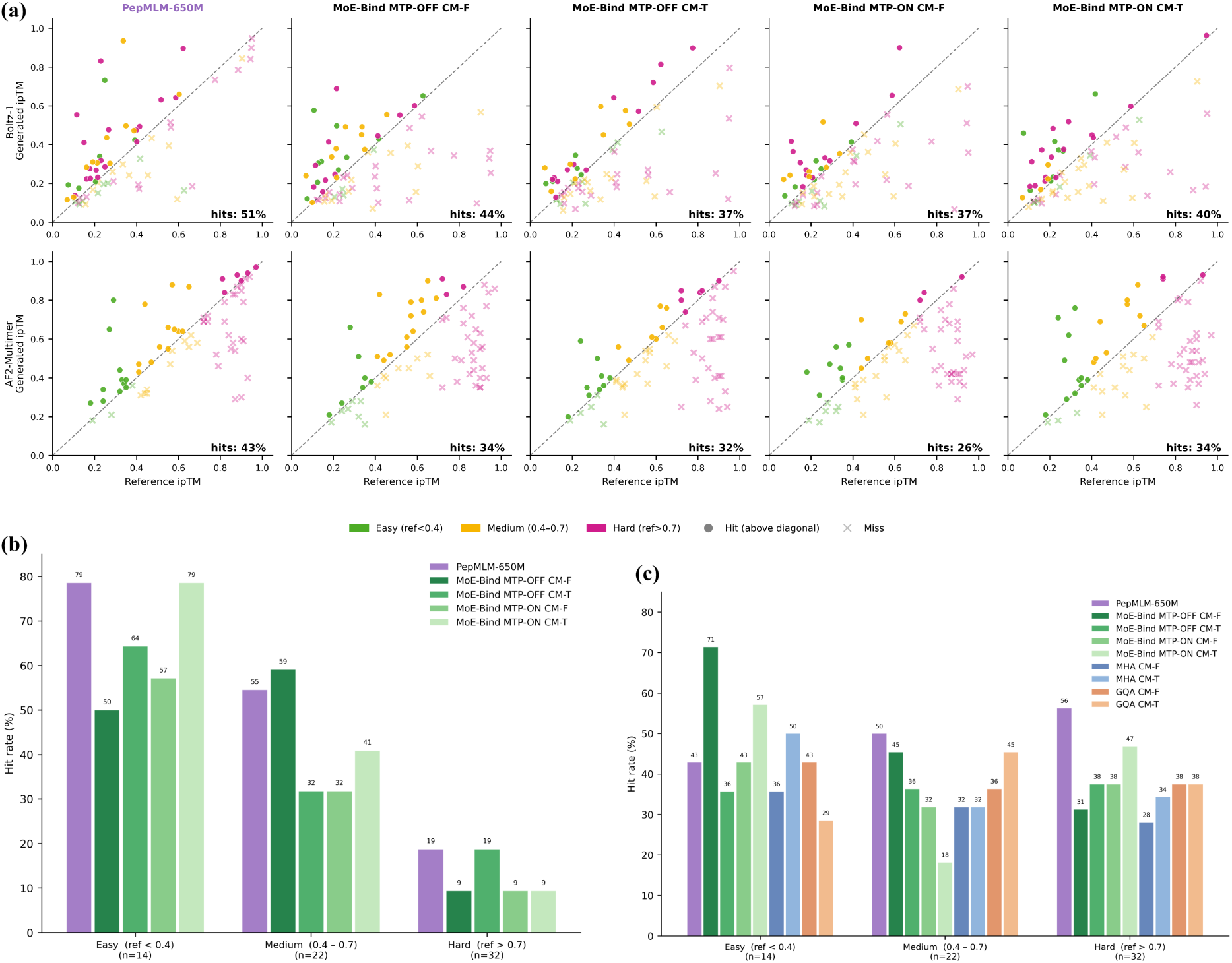
Peptide binder generation on the 68-target PepMLM benchmark, with conditional-masking (CM) and multi-token-prediction (MTP) ablations of the MoE-Bind model. **(a)** Generated versus reference ipTM for PepMLM-650M and four MoE-Bind variants, co-folded with Boltz-1 (top row) and AF2-Multimer (bottom row); points above the diagonal are hits (generated ipTM ≥ reference ipTM), coloured by target difficulty tier (Easy, reference ipTM *<* 0.4; Medium, 0.4–0.7; Hard, *>* 0.7), **(b,c)** shows per-panel hit counts. **(b)** ipTM hit rate stratified by difficulty tier using AF2-Multimer and **(c)** for Boltz-1 with reference ipTM. Abbreviations: CM F / CM T, conditional masking disabled / enabled (training loss over the full target–binder sequence vs. over binder tokens only); MTP-ON / MTP-OFF, multi-token prediction enabled / disabled during pre-training; MHA, GQA, MoE-Bind denote the three attention architectures.

**Table 7:** Aggregate peptide-binder hit rate on the 68-target PepMLM benchmark (Boltz-1, sequence-only), by architecture and conditional-masking (CM) setting. A hit is counted when the generated peptide–target ipTM meets or exceeds the native reference ipTM. Dense MHA and GQA baselines were evaluated only under Boltz-1 and have no multi-token-prediction (MTP) variant. At the matched CM-False setting, MoE-Bind (MTP-OFF) attains the highest hit rate among the architectures, exceeding both dense baselines at equal total parameters; PepMLM-650M is shown as a peptide-specialised reference.

| Architecture | Variant | CM-False |  | CM-True |  |
| --- | --- | --- | --- | --- | --- |
|  |  | Hits | Rate (%) | Hits | Rate (%) |
| MHA | dense | 21/68 | 30.88 | 25/68 | 36.76 |
| GQA | dense | 26/68 | 38.24 | 26/68 | 38.24 |
| MoE-Bind | MTP-OFF | 30/68 | **44.12** | 25/68 | 36.76 |
| MoE-Bind | MTP-ON | 25/68 | 36.76 | 27/68 | **39.71** |
| PepMLM-650M | baseline | 35/68 |  | **51.47** |  |

**Table 8:** Aggregate peptide-binder hit rate on the 68-target PepMLM benchmark under AlphaFold2-Multimer, by architecture and conditional-masking (CM) setting. A hit is counted when the generated peptide–target ipTM meets or exceeds the native reference ipTM. Only PepMLM-650M and the MoE-Bind variants were evaluated under AlphaFold2-Multimer; the dense MHA and GQA baselines were folded with Boltz-1 only (Table 7). PepMLM-650M is shown as a peptide-specialised reference.

| Architecture | Variant | CM-False |  | CM-True |  |
| --- | --- | --- | --- | --- | --- |
|  |  | Hits | Rate (%) | Hits | Rate (%) |
| MoE-Bind | MTP-OFF | 23/68 | **33.82** | 22/68 | 32.35 |
| MoE-Bind | MTP-ON | 18/68 | 26.47 | 23/68 | **33.82** |
| PepMLM-650M | baseline | 29/68 |  | **42.65** |  |

### 4.5 Routing behavior analysis of MOE in binder generation

To analyze the routing behaviour of MoE-Bind, we collected expert-routing decisions during binder generation on the 22 filtered DB5 benchmark sequences. The target generation length was set from the native ligand length; in this benchmark the ligand lengths ranged from 39 to 272 amino acids. During generation, the model was allowed to generate up to the target length plus an additional buffer, but routing statistics were computed only over the generated binder region aligned to the target ligand length. For each generated token and each MoE layer, the router selected two experts, since MoE-Bind uses top-2 expert routing. We denote the highest-probability selected expert as the first choice expert and the second selected expert as the second-choice expert. Thus, each generated amino-acid token contributes two routing assignments per layer. For routing behaviour analysis, we used the combined top-2 routing counts, i.e. both first and second choice experts were counted. For a layer *l*, expert *e*, and amino acid *a*, the plotted probability was computed as

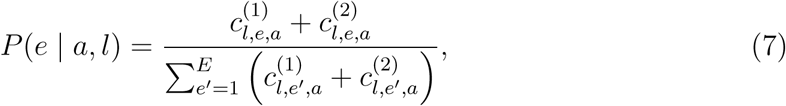

where 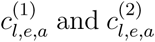 are the first- and second-choice routing counts, respectively, and *E* = 8 is the number of experts. The counts were accumulated over all 22 generated binders before normalization, so each heatmap summarizes the aggregate routing behaviour across the full benchmark. Figure 6 shows how tokens are routed across the eight experts at different layers of the model, with each column representing an amino acid and each row an expert. Brighter cells indicate that a larger fraction of tokens of that amino acid were sent to that expert. In the earliest layer (Layer 0), routing is highly concentrated, most amino acids are consistently directed to one or two dominant experts, with very little probability spread across the remaining ones. This shows that early MoE layers act as residue-specific routers, the network learns to consistently assign each amino acid to a preferred expert sub-network, suggesting that token identity alone carries sufficient signal for structured expert selection at shallow depths. This study indicates contrast to what has been observed in natural language MoE models. Jiang et al. [23] show that in Mixtral 8×7B, expert selection is nearly uniform across different text domains at all layers, with routing driven by syntactic surface form rather than meaning. The key reason for this difference is the vocabulary, natural language models work with tens of thousands of tokens where the same word can mean very different things in different contexts, making identity-based routing unreliable. Protein sequences, by contrast, have only 20 amino acids, each with fixed physicochemical properties — charge, size, hydrophobicity — that do not change with context. This gives the router a clean, stable signal to act on, enabling a level of token-level expert specialization that NLP models simply cannot achieve. Concurrent work supports this model AIDO-Protein [27], a large protein language model pretrained on 1.2 trillion amino acids using MoE, notes that experts allocate to distinct sequence patterns, however, it does not perform an explicit per-residue routing analysis or examine how this specialization evolves across layers. Similarly, recent work on expert specialization in protein MoE models [28] focuses primarily on routing of padding tokens and overall load distribution, leaving layer-wise biochemical routing patterns unexplored. To our knowledge, the present study is the first to directly visualise and interpret per-amino-acid expert routing across network depth in a protein binder generation model. Moving deeper into the network, routing progressively diffuses. By Layer 5, preferred experts still exist for many residues but probability mass begins to spread. By Layer 11, routing is distributed broadly across experts. We interpret this as a functional transition, early layers use residue identity as a direct routing signal to specialise computation, while later layers integrate long-range sequence context built up through attention, a signal that is distributed across the whole sequence and cannot be captured by looking at a single amino acid in isolation. The early-layer routing thus acts as a biochemical dispatch step that shapes how each token is processed through the network, while later experts handle the interaction-level reasoning needed for binder generation.

**Figure 6:**
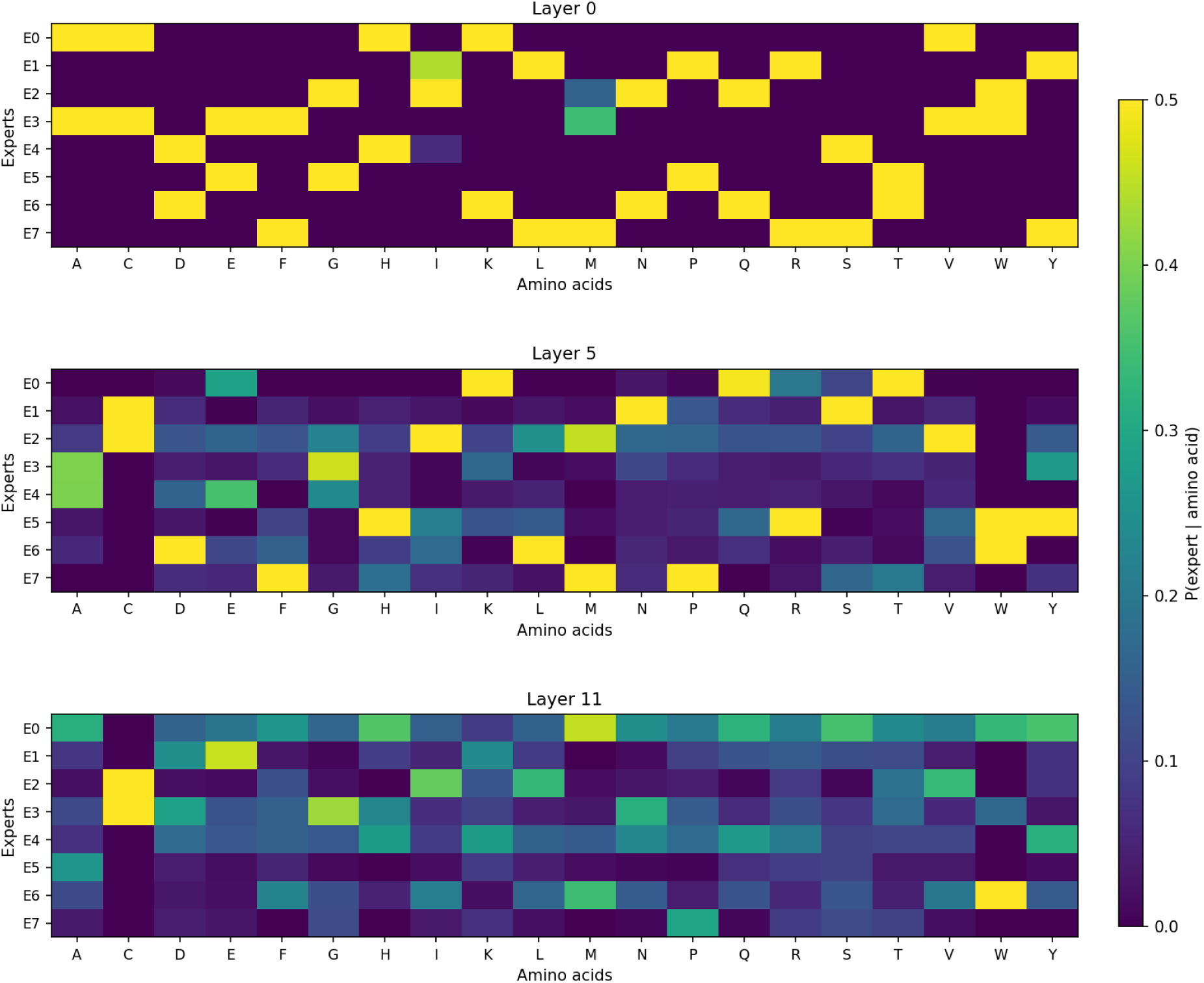
Per-layer routing behaviour of the MoE-Bind model. Each heatmap shows *P* (expert | amino acid) for a selected layer. Early layers show sparse amino-acid-dependent expert assignment, whereas deeper layers distribute routing probability across more experts, suggesting a transition from residue-identity routing to more context-aware routing.

#### 4.5.1 Expert specialization analysis of MOE

Expert specialization refers to the tendency of individual expert sub-networks within a Mixture-of-Experts model to process systematically different subsets of the input distribution. In a standard dense feed-forward layer, every token, regardless of its identity, passes through the same set of weights. MoE breaks this constraint, if the router learns to group tokens by type, each expert can develop computational routing behaviour tailored to that specific subset, increasing the effective representational capacity of the network without increasing the cost per token. For protein sequence modelling, expert specialization is particularly meaningful because amino acids fall into well-defined biochemical classes such as charged, hydrophobic, aromatic, polar etc., each of which plays a distinct structural and functional role in protein-protein interactions. If MoE routing aligns with these classes, the model is implicitly allocating separate processing circuits to biochemically distinct residue types, which is both interpretable and functionally motivated.

Figure 7 shows the routing distributions aggregated over all 12 layers and all 22 DB5 complexes. Panel (a) resolves routing at the level of individual amino acids, showing for each residue, the distribution of its first-choice (top-1) expert assignments across the eight experts. Panel (b) collapses the 20 residues into five biochemical groups and shows the usage of each expert per group using first- and second-choice routing combined, with raw token counts annotated in each cell. Panel (c) shows the first-choice enrichment ratio, how many times more frequently each expert is the primary routing target for each group compared to a uniform baseline. At the single-residue resolution of panel (a), clear amino-acid-specific preferences are already visible. The two negatively charged residues aspartate (D) and glutamate (E) both direct the largest share of their first-choice routing to E3; histidine (H) routes predominantly to E0; and isoleucine (I) routes mainly to E2. Because residues of the same biochemical class tend to share their dominant expert, these letter-level preferences aggregate cleanly into the group-level view of panels (b) and (c), which we use to characterise specialization in the remainder of the analysis. The most striking pattern is the specialization of Expert 3 (E3) for negatively charged residues (NEG; Asp, Glu). Panel (b) shows that E3 receives 25% of all combined routing events for NEG amino acids, which is nearly twice the token share expected under uniform routing and the highest value in that row. Panel (c) confirms this with a first-choice enrichment of 2.24×, the highest value in the entire matrix. At the same time, E2 almost entirely avoids NEG residues, receiving only 7% of combined NEG routing events in panel (b) and showing a first-choice enrichment of just 0.17× in panel (c) — demonstrating that specialization involves active exclusion as well as preference. E3 also shows moderate enrichment for positively charged residues (POS; Lys, Arg, His) at 1.54×, suggesting it functions broadly as a charged residue expert with a particularly strong preference for the negative charge class. E6 acts as a secondary expert for NEG at 1.71×, further reinforcing the concentration of negatively charged residue processing within a small subset of experts.

**Figure 7:**
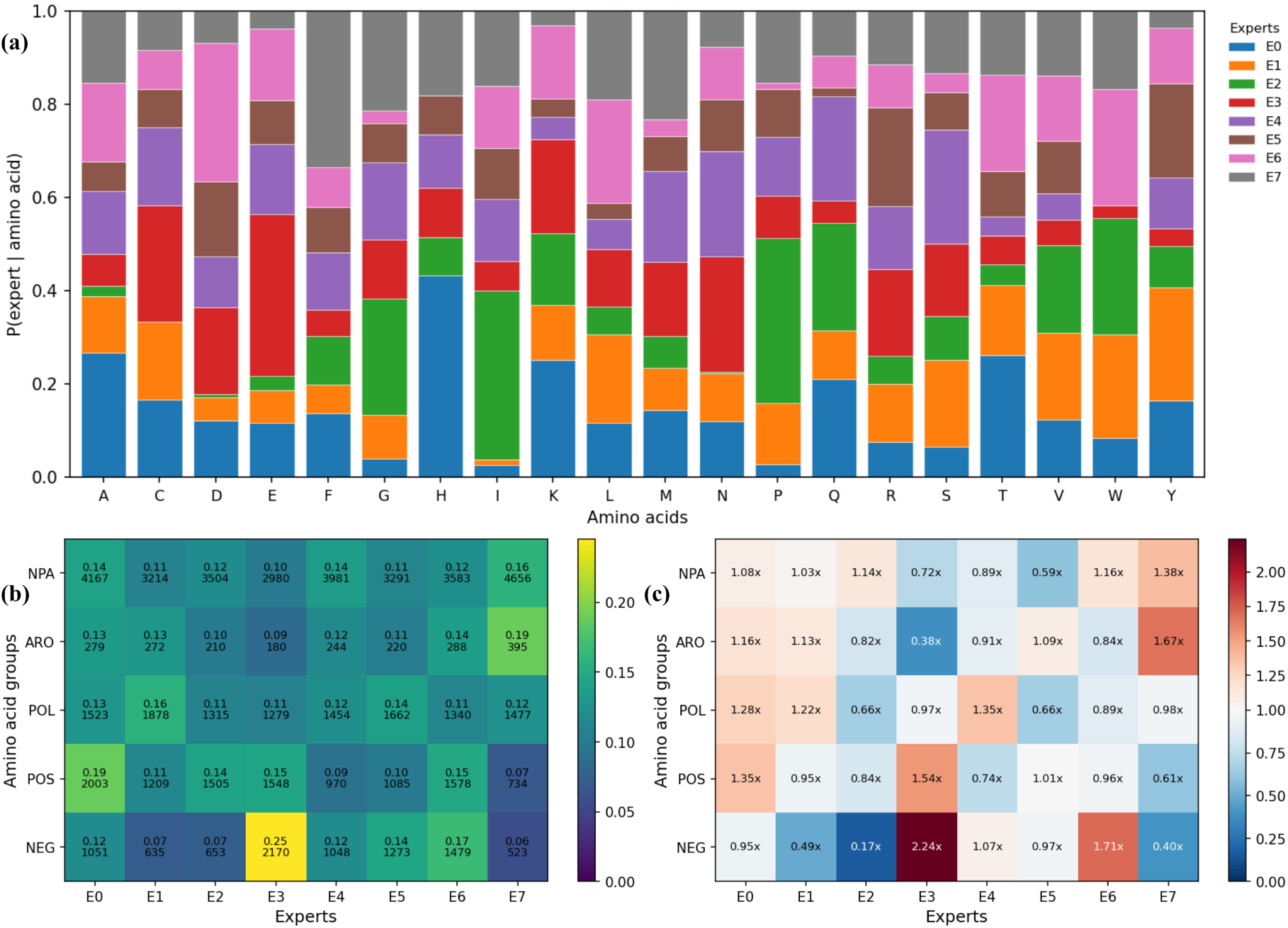
Expert routing specialization by amino acid, aggregated over all 12 MoE layers and 22 DB5 benchmark complexes for the MoE-Bind. (a) First-choice (top-1) routing resolved per individual amino acid; each bar sums to one over the eight experts. (b) Expert usage across the five amino acid biochemical groups using first- and second-choice routing combined; raw token counts are annotated below each probability value. (c) First-choice enrichment ratio relative to uniform routing (1.0×); red indicates over-representation, blue indicates under-representation.

A second clear pattern emerges for hydrophobic residues. Expert 7 (E7) shows the highest enrichment for both aromatic residues (ARO; Phe, Tyr, Trp) at 1.67× and nonpolar aliphatic residues (NPA; Gly, Ala, Val, Leu, Ile, Met) at 1.38× in panel (c). Panel (b) confirms that 19% of aromatic routing events go to E7 — the highest value in the ARO row, and 16% of nonpolar aliphatic events, the highest in the NPA row. This positions E7 as a hydrophobic residue expert handling both bulky aromatic side chains and the smaller aliphatic residues that form the hydrophobic core of protein structures. Notably, E3 shows near-complete avoidance of aromatic residues (0.38×), reinforcing that E3 and E7 operate on complementary biochemical subspaces — one specializing in charge, the other in hydrophobicity.

The remaining experts show subtler but consistent preferences. For positively charged residues, E0 receives the highest combined fraction in panel (b) at 19% with a 1.35× first-choice enrichment, while E3 contributes an additional 1.54×; this is consistent with the strong His with E0 preference already visible at the single-residue level in panel (a). For polar uncharged residues (POL), E1 is the most preferred expert at 16% in panel (b) and 1.22× in panel (c), followed by E0 at 1.28×. The nonpolar aliphatic group, being the largest biochemical class by token count, is distributed most broadly across experts, reflecting both the internal diversity of this group and the load-balancing pressure that prevents any single expert from monopolising too large a share of tokens. These preferences are visible in the all-layer aggregated view indicates that routing specialization is a consistent and stable property of the network across depth, not a feature of any single layer. The structured assignment of biochemical classes to dedicated experts shows that the MoE architecture, even at the small scale, develops an implicit functional partitioning of amino acid space, that emerges from training signal alone without any explicit supervision toward biochemical grouping.

### 4.6 Computational Efficiency of the MoE-Bind Architecture

The MoE-Bind (100M) architecture achieves substantial computational savings over both the MHA (100M) and GQA (100M) baselines through two complementary mechanisms, sparse expert activation and low-rank KV compression. Although all three architectures occupy a comparable storage footprint (∼100 M total parameters), MoE-Bind activates only 38.8 M parameters per token versus 99.85 M (MHA) and 100.74 M (GQA), a 2.57× reduction in active compute. Because training cost scales with active, not total, parameters, this translates directly to a 2.57× reduction in estimated floating-point operations, 78 M FLOPs per token for MoE-Bind compared with 199.7 M (MHA) and 201.5 M (GQA).

Accumulated over the full pre-training and fine-tuning budgets, MoE-Bind consumed 354,662 TFLOPs in total of a 56.2% reduction relative to MHA (810,526 TFLOPs) and a 57.7% reduction relative to GQA (838,135 TFLOPs), as measured by hardware FLOP counters during training (Table 9). MHA and GQA are nearly identical in training cost because neither employs sparse activation, all parameters participate in every forward pass, and reducing the number of KV heads in GQA, while beneficial for inference memory, does not meaningfully reduce the total parameter count or per-token FLOPs at this scale, where MLP layers dominate training compute. Standard MHA caches full key and value tensors for every attention head at every layer. At sequence length 768 with batch size 1 in fp16, this amounts to 32,256 KB for MHA, GQA reduces this to 12,288 KB (2.6× reduction) by sharing KV projections across groups of query heads, MLA compresses further by projecting keys and values into a low-rank latent space before attention, caching only this compressed representation rather than the full per-head tensors. The resulting KV-cache is 1,152 KB, a 28.0× reduction relative to MHA and a 10.7× reduction relative to GQA (Figure 8a). This is particularly consequential for protein-sequence generation, where context lengths can be large based of the sequence length; a smaller KV-cache allows larger batches or longer sequences without increasing peak GPU memory.This is particularly relevant for high-throughput binder screening, where many receptor-binder pairs must be evaluated in parallel. At small batch sizes, model weights dominate memory and the KV-cache difference is negligible, however, as batch size increases the KV-cache becomes the dominant memory consumer for MHA and GQA, whereas MoE-Bind’s compressed representation remains a minor contributor, allowing proportionally larger batches within the same GPU memory budget. Figure 8b shows that MoE-Bind maintains a total parameter count of 102.7 M with equivalent of the dense baselines, while activating only 38.9 M parameters per token, comparable to the 38 M-parameter dense models trained in a separate compute-matched tier. This occupies a unique position, MoE-Bind is storage matched to the 100 M-scale dense models yet compute matched to the 38 M-scale dense models, enabling a two-way comparison within a single trained checkpoint. During inference due to MoE routing and MLA decompression overhead, single-sequence generation throughput is lower than the dense baselines at batch size 1. However, this trade-off is offset by the consistently stronger binder generation results observed for MoE-Bind across evaluations, alongside a 56.2% reduction in training FLOPs and a 28× smaller KV-cache footprint demonstrating that MoE-Bind delivers superior sequence quality at substantially lower computational and memory cost than either dense counterpart.

**Figure 8:**
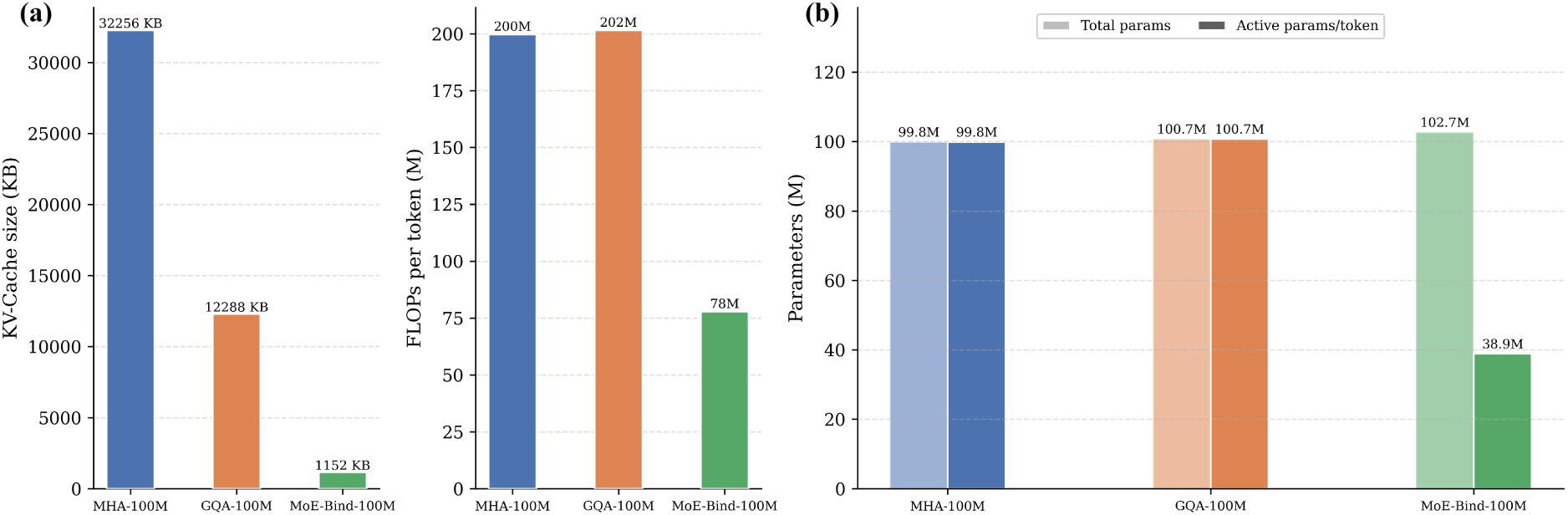
Theoretical inference-time efficiency across architectures. (a) KV-cache memory and FLOPs per token at sequence length 768, batch size 1, fp16. (b) Total versus active parameters per token.

**Table 9:** Cumulative training compute (TFLOPs) measured across pre-training and finetuning for each architecture, with MoE-Bind savings relative to MHA and GQA. Values are hardware FLOP counts logged during training.

| Stage | MoE-Bind<br>(TFLOPs) | Baselines |  | MoE-Bind Savings |  |
| --- | --- | --- | --- | --- | --- |
|  |  | MHA (TFLOPs) | GQA (TFLOPs) | vs. MHA (%) | vs. GQA (%) |
| Pre-training | 233,149 | 532,854 | 550,975 | 56.2 | 57.7 |
| Fine-tuning | 121,513 | 277,672 | 287,161 | 56.2 | 57.7 |
| **Total** | **354,662** | **810,526** | **838,135** | **56.2** | **57.7** |

## 5 Discussion

The most consistent observation across our experiments is that the MoE-Bind architecture is able to match or exceed compute-matched dense baselines while activating less than half their per-token parameters. We interpret this not as a generic confirmation that sparse models help, which has been established amply in natural language, but as a statement about what binder generation actually requires. Receptor-conditioned binder generation is a task in which different residue positions impose qualitatively different constraints, hydrophobic core packing in one region, charged interface complementarity in another, hinge flexibility elsewhere. A dense feed-forward stack must answer all of these constraints through a single shared parameter set, which forces representational compromises across positions. A sparse mixture-of-experts stack instead allows the router to send each residue position to the sub-network whose learned parameters are most relevant for the local biochemical context, so the same total budget is spent more selectively. In natural language, expert specialization in MoE models is usually diffuse and hard to map onto interpretable token categories, because the vocabulary spans tens of thousands of sub-word units and any regularity is spread thinly across many tokens. Proteins are different in the sense that, effective token alphabet is fixed at only twenty residues, and these residues fall into a small number of well-known biochemical groups such as hydrophobic, polar, charged, and aromatic. This narrow, biologically structured vocabulary acts as a built-in inductive prior on what kinds of specialization can plausibly emerge in the first place. The expert assignments we observe match this prior closely, with dedicated experts for charged and aromatic residues and broader sharing across the larger aliphatic class, without any explicit supervision toward biochemical grouping.This suggests that MoE-Bind’s experts are partitioning the protein vocabulary along the same biochemical lines that biology already recognises, and that proteins may turn out to be a particularly clear and natural domain for studying interpretable expert routing in sparse generative models.

The contribution of Multi-head Latent Attention in our experiments complements that of the MoE feed-forward stack. Where MoE lowers the active compute spent per residue, MLA lowers the active compute spent inside the attention layer itself. Standard attention stores a separate set of keys and values for every attention head at every layer, which inflates both the training cost and the per-token compute at inference. MLA replaces this with a low-rank latent projection, so the model only learns and updates a small compressed representation rather than the full per-head tensors. This compression carries through to both training and generation. During training it reduces the parameter count and the per-step FLOPs of the attention block, contributing directly to the lower training compute reported in our efficiency experiments. During generation it reduces the work performed at each decoding step, so MoE-Bind produces each new residue with substantially less active compute than its dense peers, while still preserving the per-head expressivity that binder generation appears to depend on at the interface. Long-receptor caching benefits naturally from the same compression, but the primary benefit is the consistent active-compute reduction that MLA contributes alongside MoE across the entire pipeline. State-of-the-art structure-based binder design systems such as RFdiffusion, Al-phaProteo, BindCraft, and BoltzGen remain the methods of choice when high-resolution target structures and substantial compute budgets are available, and continue to deliver binders with high success rates. MoE-Bind operates in a different regime and addresses a different need. At only 100M parameters, with no structural input during generation, and at a fraction of the active compute of its dense counterparts, MoE-Bind produces binders whose interface confidence matches or exceeds parameter-equivalent dense sequence-only baselines and approaches a peptide-specialised reference model on a benchmark for which it was never specifically trained. This makes MoE-Bind a strong complement to structure-based pipelines rather than a competitor. In an upstream role, it can rapidly generate and intrinsically filter hundreds to thousands of candidate binders per target, producing a small high-quality shortlist that structure-based systems can then refine and rank under their usual folding and inverse-folding steps; this shifts the expensive structural computation from every candidate to only the surviving few. For the very large class of targets without a usable three-dimensional structure, sequence-only generation is the only practical route to de novo binder design, and MoE-Bind is purpose-built for this regime. It is also attractive when compute is constrained or when many targets must be explored in parallel.

MoE-Bind is trained at a deliberately modest scale, 100M total parameters and around 39M active per token, on a compute budget consistent with the resource constraints under which the work was carried out. This scale is enough to isolate the architectural effects of MoE-Bind under a compute-fair comparison against dense baselines, and the resulting performance is favourable against parameter-equivalent dense models. The upper bound of what the MoE-Bind architecture can ultimately reach would only become visible at the billion-parameter scale that characterizes state-of-the-art natural-language MoE systems, which remains an open direction for future scaling work. The structural evaluation also reflects the well-documented fact that state-of-the-art predictors such as Boltz-2 and AlphaFold2-Multimer can disagree on a per-complex basis, which is one reason we report results under both predictors; the broad agreement between them across the benchmark is what we treat as the load-bearing signal. A natural extension is to include further independent predictors as they mature, and to quantify per-complex inter-predictor agreement as an additional confidence layer on top of the ipTM hit criterion. The routing analysis is consistent across layers and across the DB5 benchmark, but it is based on a single 100M checkpoint with eight experts. A natural next direction is to move from observed specialization to deliberate specialization. Because individual experts in MoE-Bind already align with specific amino acids and biochemical groups without supervision, a lightweight auxiliary objective could explicitly assign each expert to a predefined biochemical class, residue identity, or even higher-order motif such as charged interface patches or hydrophobic core packing. Once experts are aligned in this way, the router becomes a user-facing control surface. A user could bias generation toward binders enriched in particular residues or biochemical compositions by up-weighting the corresponding experts at inference time, producing binders tailored to the chemical or functional profile required for a given target. This would convert the interpretable routing reported here into a practical mechanism for controllable sequence-only binder design, and we view it as one of the most promising directions opened by the present work.

Taken together, these results show that sparse Mixture-of-Experts and Multi-head Latent Attention transfer cleanly into target-conditioned protein binder generation, with selective per-residue specialization in the feed-forward stack and active-compute compression in the attention stack working together to deliver strong sequence-based binders. MoE-Bind establishes that this combination of architectural choices is the right direction for de novo protein binder design. In natural language, autoregressive decoder based models remain the dominant paradigm and continue to outperform every competing architecture as they are scaled and refined, and the same trajectory is plausible for protein binder generation, where autoregressive sequence modelling has so far been explored at only a small fraction of the scale and architectural maturity of that natural language. With proper architectural tuning, scale, and integration with structure-based refinement, MoE-Bind opens a clear path toward this potential, and toward translating sequence-only binder generation into biological function at the level of real therapeutic targets.

## Data and Code Availability

The code is open-sourced via GitHub and is available at https://github.com/Dipayan26/ MoE-Bind. All datasets used in this study are derived from publicly available resources.

## Author Contributions

**Dipayan Sarkar:** Conceptualization, Methodology, Software, Investigation, Formal analysis, Data curation, Visualization, Writing – original draft, Writing – review and editing. **Chiranjib Sarkar:** Conceptualization, Supervision, Project administration, Resources, Writing – review and editing.

## Declaration of Generative AI

During the preparation of this work the authors used Anthropic’s Claude solely for language refinement; all experiments, analyses, results, figures, citations, and scientific interpretations are the authors’ own, and the authors reviewed every passage and take full responsibility for the content of the publication.

## Acknowledgements

The authors thank the Department of Bioinformatics, University of North Bengal for providing computational resources and infrastructure that supported this work. We thank India AI Compute initiative (Project ID: P1-S2025070964) for providing us compute infrastructure. Dipayan Sarkar acknowledges the University Grants Commission (UGC), Government of India, for financial support through the CSIR-UGC Junior Research Fellowship.

